# Flight-tone mediated circadian control of audibility in mating swarms of *Anopheles* mosquitoes

**DOI:** 10.1101/2021.07.12.452033

**Authors:** Jason Somers, Marcos Georgiades, Matthew P Su, Judit Bagi, Marta Andrés, Gordon Mills, Watson Ntabaliba, Sarah J Moore, Roberta Spaccapelo, Joerg T Albert

## Abstract

Mating swarms of malaria mosquitoes form every day at sunset throughout the tropical world, they typically last less than 30 minutes. Activity patterns must thus be highly synchronized between the sexes. Moreover, males must be able to identify the few sporadically entering females by detecting the females’ faint flight tones. We here show that the *Anopheles* circadian clock ensures a tight synchrony of male and female activity and –importantly – also retunes the males’ acoustic detection system: by lifting their own flight tones at dusk, males actively enhance the audibility of females. The reported phenomenon of ‘harmonic convergence’ is a random by-product of the mosquitoes’ flight tone variance. Intriguingly, flight tones of individual mosquitoes occupy narrow –partly non-overlapping-frequency ranges, suggesting that the audibility of individual females varies across males.

**One Sentence Summary:** Male mosquitoes sharpen their hearing at sunset.

---

In sexually reproducing animals two mating types, or sexes (most commonly males and females), must find each other for the act of copulation. Yet both extrinsic (e.g. environmental) and intrinsic (e.g. sexual-dimorphism-related) factors can lead to asymmetries in the spatial and temporal dispersal of the sexes (*1*). This is also true for disease-transmitting mosquitoes (*2*).

Mosquitoes display numerous sexually dimorphic traits, including the female-specific blood-feeding behavior (*3*) – and the male-specific daily formation of mating swarms (*4*). Mating swarms, however, are also part of the solution to the dispersal problem: hundreds or thousands (*5*) of males congregate at a fixed location, guided by visual markers (*6*), and at a fixed daytime, typically dusk (*7*), to act as reproductive mates for a much smaller number (a few dozens) of sporadically entering females.

In the malaria-vector species of the *Anopheles gambiae* complex, mating swarms form a crucial reproductive bottleneck, making them a prime target of current vector control efforts (*7*). While *Anopheles* mating swarms can form reliably at the same sites - and same daytimes - for years on end, individual swarm durations of sometimes less than 20 min. (*8, 9*) also make them an astonishingly ephemeral phenomenon. The short-lived nature of the swarms together with the sparsity of females are intriguing from two scientific points of view. Chronobiologically, it implies a tight synchronization between male and female activities. Neurobiologically, it suggests a highly efficient operation of the sensory systems that guide the males’ mating behavior. A key sensory modality for a male’s copulatory success is his sense of hearing (*10*). In mosquitoes, copulae between male and females form in mid-air and are preceded by an acoustic chase, where a male follows the flight tone of a female. This long-known male behavior (*11*), called phonotaxis, must succeed against the backdrop of the flight tones of hundreds of other males and constitutes one of the most reproducible, and most impressive, behaviors in insects.

Mosquito hearing relies on an active process (*12*); the flagellar receivers pick-up air-borne vibrations (*13–15*), which are transduced into electrical currents – and mechanically amplified - by mechanosensory neurons of Johnston’s Organ (JO) (*11*). The JO of an *Anopheles gambiae* male is exquisitely sensitive and responds to flagellar tip deflections of <20nm (or <1mdeg, respectively) (*16*). For a mosquito, however, hearing goes beyond the simple *reception* of external sounds as it also involves – and partly necessitates - the *generation* of sound. The mechanistic explanation for these settings lies in the way mosquito hearing works (*17*). The operation of the mosquito ear introduces essential nonlinearities (e.g. gating compliances (*16*)) into the mechanics of its flagellar sound receiver. As a result of its nonlinearities, a stimulation with two pure tones will generate additional, mathematically predictable distortion products (DPs) (*18*) to the receiver’s motion. For the ear of a flying male, some of the lower-frequency DPs that are generated by the mixing of his own flight tone with the flight tone of a near-by flying female will be more audible than the actual flight tones themselves (which are mostly inaudible) (*19*). Hearing - or more broadly audibility - in mosquitoes is thus inextricably linked to their flight activity and in fact dependent on a specific interrelation between male and female flight tones and the distortions these produce (*20*).

One such relational state - described as ‘harmonic convergence’ (HC) (*21*) - has been interpreted as acoustic interaction between males and females (*22*). To investigate the relationships between HC, DPs and the acoustic environment within swarms, we surveyed the daily flight tone landscape of *Anopheles gambiae*, specifically probing for circadian modulations of audibility related to the mating swarm (*23*). We also quantified the relative contributions made by males and females, respectively.

## Results

### Tight circadian synchrony between the sexes

To probe behavioral synchronicity between the sexes, we monitored baseline activities (1 min. binning, LAM25H-3, Trikinetics) separately in *Anopheles gambiae* males and females. Both sexes were exposed to the same environmental sequence: 4 initial days of LD entrainment (consisting of 11h-long days and nights, flanked by 1h-long artificial ‘sunrises’ and ‘sunsets’, at 28°C and 80%RH) were followed by 5 days without any temporal cues (‘free-running conditions’: complete darkness, at 28°C and 80%RH).

Under LD conditions, the activity patterns of males and females were highly similar (Fig. 1): an initial ‘lights-on’ startle response at ‘sunrise’ was followed by a near complete inactivity during the rest of the day (Fig. 1A). Main activities were shown at ‘sunset’. Males started their activity increases earlier, and maintained them for longer, than females (Fig. 1B). Both sexes responded to lights-off (at ZT13 and coinciding with an illuminance drop from ∼20 lx to zero) with an immediate and steep activity increase. The males’ activity plateau lasted for <30min and fully enveloped the female activity peak (Figure 1B). During entrainment, the males’ and the females peak activities differed by ∼1min (Fig. 1C). Even in the absence of external temporal cues, male and female activities remained tightly synchronized. On the 5^th^ day of free-running conditions the time difference between male and female peak activities was only 16.93min±35.12min SEM (p-value = 0.63, females = 20, males = 26) and the 5-day average difference between the sexes was less than 30 min (23min±12.47min SEM, p-value 0.14, n = 5) (Figure 1C). The synchrony between the sexes persisted although their circadian clocks ran considerably faster than 24h (free running period in males: 22.64h±0.12h SEM; females: 22.54h± 0.11h SEM; Fig. S1) and thus activity peaks themselves constantly shifted to earlier daytimes (Fig. 1C, bottom).

**Fig. 1.**
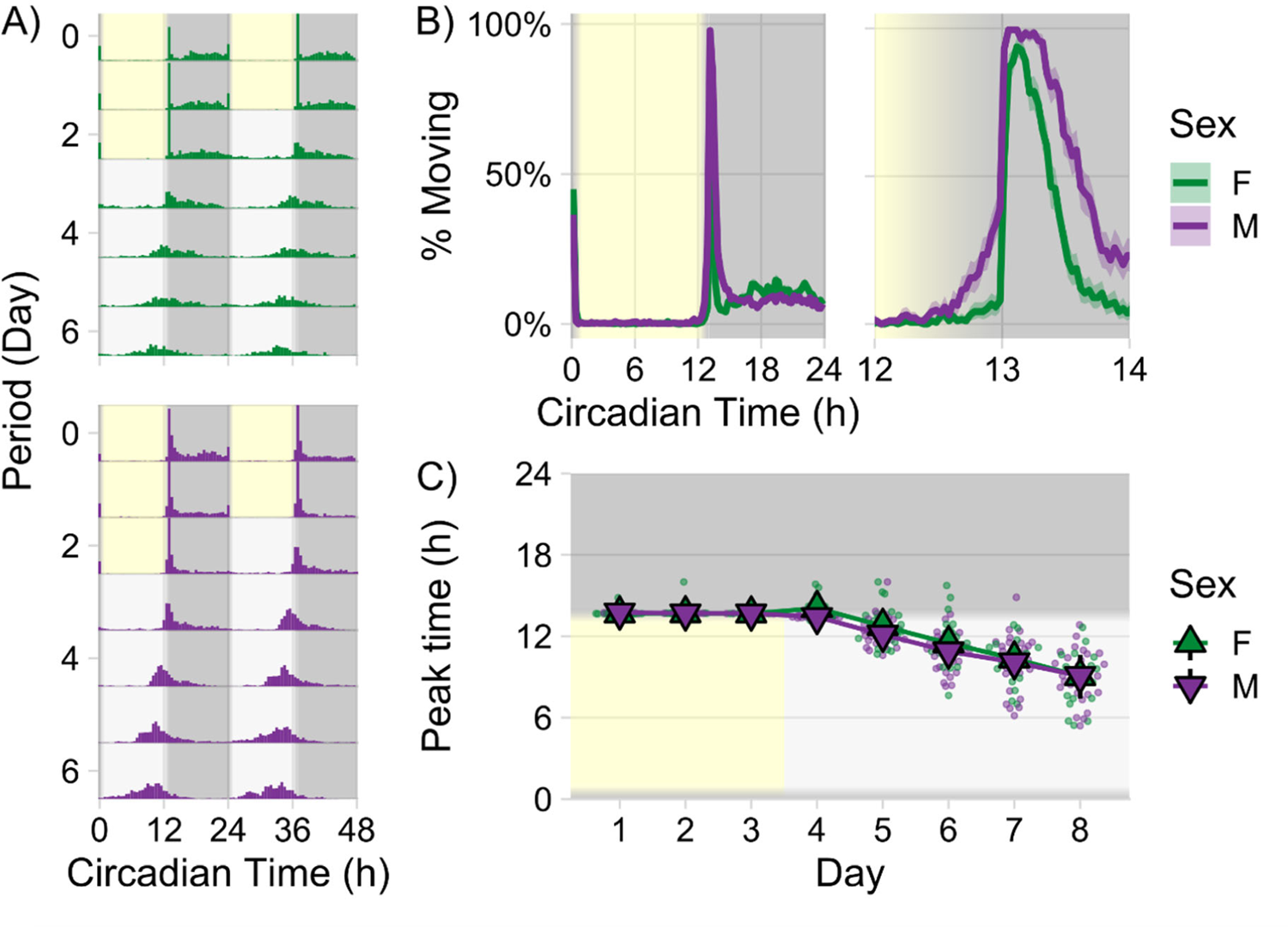
Circadian locomotor analysis of mosquitoes held in activity monitors (Trikinetics) A) Double-plotted actograms for entire experiment - 3 days of 12-hour light/dark entrainment (with 1 hour dusk/dawn simulations) at 28°C followed by 5 days of free-running conditions (constant darkness, constant temperature). Activity data is binned at 30 minutes and plotted in overlapping 48-hour intervals to visualize periodicity. B) Average locomotor activity across 3 entrainment days (left: for entire 24 hours, right: for 2 hours around dusk). Activity is plotted as percentage of groups moving ± S.E.M. C) Daily peak activity times across the entire experiment, calculated as time of highest activity from low-pass filtered raw data (>30 mins). Triangles display group medians ± 95% C.I. (n = 20 females and 27 males). Data is shown from 1 individual experiment.

### Male-driven audibility boost in swarming *Anopheles*

A very close temporal alignment was also seen in the activity peaks of free-flying populations of *Anopheles* males and females (100-strong each). When assessed in separate cages, male and female activity peaks (quantified acoustically by the number of flyby events past a stationary microphone, Fig. 2A,B) differed by only 1.5min ± 0.25min S.E.M in LD and by 1.75min ± 3.5min S.E.M. in the first day of free-running conditions (Fig. S2).

**Fig. 2.**
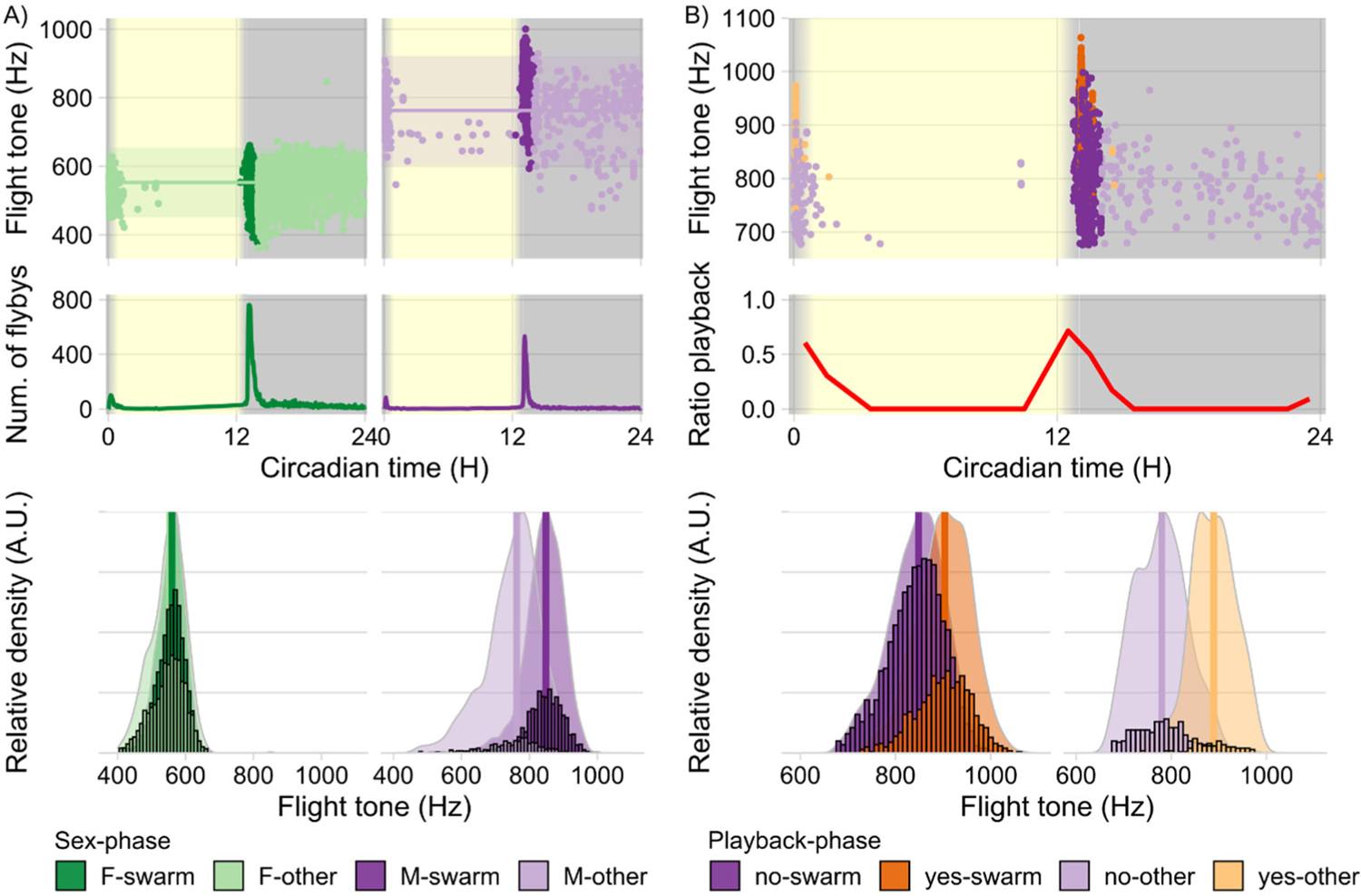
Circadian acoustic analysis of free-flying populations of male and female *Anopheles* mosquitoes. A) *Top panels* Individual flight tones recorded from free-flying populations of females (green) and males (purple) during 12-hour light/dark entrainment (with 1 hour simulated dusk and dawn) at 28°C. Points represent median flight tones of individual flyby events (see online Methods), darker colors indicate swarm time. Solid horizontal lines show the population medians for out-of-swarm flight tones ± 95% C.I. for each sex (F = 552.96 Hz, σ = 50.70 Hz, n = 2339 flight tones; M = 762.67 Hz, σ = 79.95 Hz, n = 445 flight tones). *Middle panels* Line plots showing running averages (window: 5 min) of number of recorded flight tones across the day as a measure of flight activity. Mean peak activity times across all entrainment days was calculated for each sex (F = CT 13.18 H, σ = 0.01 H, n = 12 days; M = CT 13.15 H, σ = 0.01 H, n = 18 days). *Bottom panels* Distribution of flight tones recorded for each sex across the entrainment days separated by phase – swarm and out-of-swarm (other). Bar plots are binned counts of the individual flight tones and they are plotted against scaled density plots in order to visualize the distribution shift between the swarm and out-of-swarm (other) group. Vertical lines indicate the calculated median for each sex-phase combination (F-swarm = 559.95 Hz, σ = 40.42 Hz, n = 3040 flight tones; F-other = 552.96 Hz, σ = 50.70 Hz, n = 2339 flight tones; M-swarm = 848.85 Hz, σ = 55.30 Hz, n = 1600 flight tones; M-other = 762.67 Hz, σ = 79.95 Hz, n = 445 flight tones). B) *Top panels* Individual flight tones recorded from populations of free-flying male mosquitoes presented with a 1-minute artificial female flight tone (550Hz) at 30-minute intervals. *Middle panels* Line plot shows ratio between flight tones recorded during playback of artificial female flight tone and the total number of flight tones within each 30-minute interval (n_(playback)_/n_(total)_). *Bottom panels* Bar plots are binned counts of individual flight tones; they are plotted against scaled density plots in order to visualize the distribution shift between the playback and no playback group for each experimental phase. Vertical lines indicate the calculated median for each playback-phase combination (no-swarm = 848.20 Hz, σ = 58.25 Hz, n = 2318 flight tones; yes-swarm = 903.48 Hz, σ = 57.67 Hz, n = 928 flight tones; no-other = 780.04 Hz, σ = 53.80 Hz, n = 354 flight tones; yes-other = 888.96 Hz, σ = 47.77 Hz, n = 92 flight tones). Female data are pooled from 3 independent experiments, male data are pooled from 4 independent experiments and playback data are pooled from 4 independent experiments.

More intriguingly though, male – but not female - flight tones varied significantly across the day (Fig. 2A,B). At 28°C, and under LD conditions, male flight tones had a median frequency of 849 ±55 Hz during swarm time (±30 min around the circadian sunset at ZT13) as compared to all other times of the day (762 ±80 Hz) (Fig. 2A, right). Female flight tones did not show significant daily variations (Fig. 2A, left). The corresponding ratio between male (*f_2_*) and female (*f_1_*) flight tones was 1.38 ± 0.20 for most of the day, but the male-specific increase lifted it to a value of 1.53 ± 0.16 around sunset.

In line with their daily flight activity patterns, one-hundred-strong populations of male *Anopheles* showed phonotactic responses (measured as increases of flyby events past the source of a 550Hz pure tone) predominantly at swarm time, i.e. at sunset (Fig. 2B, top). Some, albeit reduced, phonotactic responsiveness was also seen in the morning (at sunrise) but responses were virtually absent throughout the rest of the day (Fig. 2B, middle).

After playback of a female-like tone (550Hz), the median flight tones of male *Anopheles* rose further to 903 ±58 Hz during swarm time (Fig. 2B, bottom left). This *phonoacoustic* response lifted the male/female flight tone ratio to 1.62 ± 0.2. The flyby increases shown by *Anopheles* males after playback at other times of the day also associated with an upshift of their own flight tone frequencies (Fig. 2B, bottom right), suggesting a close link between phonotactic and phonoacoustic response. The key behavioral changes observed around ZT13 - i.e. the increases in flight activity and flight tones - persisted under free-running conditions, demonstrating their circadian origin (Fig. S3).

These findings show that the flight tones of male *Anopheles* can assume distinct frequency states. For the vast part of the day they occupy a *baseline state* characterized by low frequencies; around sunset, flight tones increase in frequency and move to a *swarming state*; after detection of a female-like flight tone they finally raise their frequencies even higher and enter an *activated state*. The finding that these three states are centered on a male/female flight tone ratio of 1.5 (baseline: 1.38, swarming: 1.53, activated: 1.62) is interesting and relevant. For a two-tone, and distortion-based, hearing system like that of mosquitoes, an interval ratio of 1.5 (also called the perfect fifth in music theory) constitutes a singularity. This results from two low-frequency DPs, the quadratic DP (or ‘difference tone’), *f*_2_*-f*_1_, and the cubic distortion product, *2f*_1_*-f*_2_ becoming numerically identical at a (*f*_2_*:f*_1_) ratio of 1.5, thus creating a ‘super distortion’ (Fig. 3A).

**Fig. 3.**
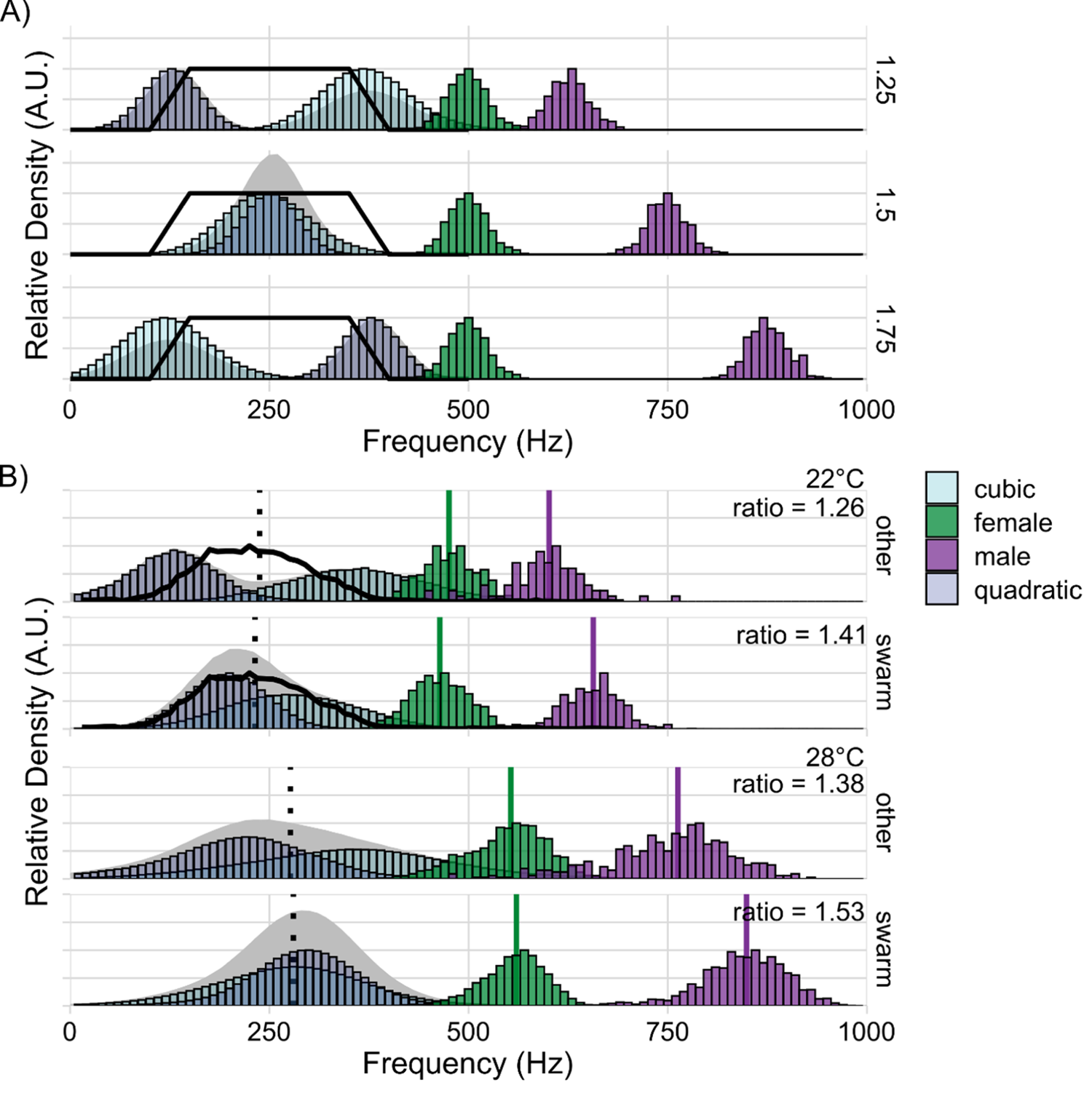
Flight-tone controlled generation of audible distortion. A) Conceptual plot of the quadratic (f2 – f1) and cubic (2*f1 – f2) distortion tones produced by the non-linear mixing of female (f1) and male (f2) flight tones. Distortion tones were calculated for hypothetical populations of one female (μ = 500 Hz, σ = 25 Hz, n = 500 flight tones) and three male flight tones (μ = 625; 750; 875 Hz, σ = 25 Hz, n = 500 flight tones) to demonstrate the respective distortion tone distributions for different male:female flight tone ratios. The solid black line creates a hypothetical window of the spectral sensitivity of the male auditory nerve. At population mean ratio of 1.5 a singularity occurs, quadratic and cubic distortion tones become numerically identical and superimpose at a frequency of 0.5*f1. B) The same distortion tones were calculated with experimental data to compare swarm and out-of-swarm (other) distributions at 22°C. Here, we could also match our own nerve recordings (solid black lines in 22°C data) to the created distortions. The vertical, dotted black line demarcates an optimum prediction for the center of male nerve responses for a distortion product based hearing system (optimum frequency = 0.5*f1). At both 22°C and 28°C the overlap between quadratic and cubic distortion tones is greater at swarm time than at out-of-swarm times. The colored vertical lines indicate the calculated median flight tone for each sex (22°C/other: females = 475.39 Hz, σ = 36.85 Hz, n = 447 flight tones; males = 601.08 Hz, σ = 51.97 Hz, n = 83 flight tones; 22°C/swarm: females = 464.03 Hz, σ = 37.04 Hz, n = 1137 flight tones; males = 656.53 Hz, σ = 39.01 Hz, n = 454 flight tones; 28°C/other: females = 552.96 Hz, σ = 50.70 Hz, n = 2339 flight tones; males = 762.67 Hz, σ = 79.94 Hz, n = 445 flight tones; 28°C/swarm: females = 559.95 Hz, σ = 40.41 Hz, n = 3040 flight tones; males = 848.85 Hz, σ = 55.30 Hz, n = 1600 flight tones). Average ratios are calculated for every combination of male and female flight tone for each temperature/time combination (22°C/other = 1.26, σ = 0.15, n = 37,101 pairs; 22°C/swarm = 1.41, σ = 0.14, n = 516,198 pairs; 28°C/other = 1.38 σ = 0.20, n = 1,040,855 pairs; 28°C/swarm = 1.53, σ = 0.16, n = 4,864,000 pairs). Both female and male 22°C data are pooled from 2 independent experiments..

Mosquito flight tones have been reported to vary with ambient temperature (*24*). We tested the daily distribution of flight tones also at 22°C and observed the same phenomena. At swarm time, the flight tone ratio was 1.41 ± 0.14 but dropped to 1.26 ± 0.15 for the rest of the day (Fig. 3B) and - as was the case at 28°C - flight tone changes were restricted to males.

At 22°C we could now also directly compare the frequency response function (resolution Δf=15Hz) of the males’ nerves to the spectral bandwidth of DPs generated by the observed flight tones. In all ears tested, compound nerve responses to sinusoidal flagellar oscillations were limited to frequencies between ∼65Hz and ∼400Hz, with a plateau of maximal responses occurring between ∼150Hz and ∼300Hz (Fig. 3B and Fig. S5). Male ears can assume two states, a quiescent (baseline) state and a state of self-sustained oscillations (SSOs) (*16*). In the baseline state, no ear tested showed any response to frequencies >450Hz (Fig. S5). Higher frequency responses have been reported though (*21, 25–27*), and also occurred in our recordings, all of these, however, were linked to SSOs. In the absence of another tonal component (a second tone or an SSO), which contributes to the production of audible distortion, no response to higher frequencies occurred (data not part of this manuscript, but available on request). Comparing the sensitivity of male *Anopheles* nerves to the DPs available to their ears at swarm time, with those DPs occurring during the rest of the day, shows how the male-specific flight tone increase harvests female audibility by increasing the overlap between cubic and quadratic DPs (Fig. 3B).

### ‘Harmonic convergence’: an epiphenomenon of flight tone variance

Our data show how daily modulations of flight tone changes in male *Anopheles* adjust, and optimize, the audibility of females specifically at swarming time. These modulations are centered on a male/female flight tone ratio of 1.5, which also constitutes an important theoretical optimum. All our data was gathered from separately kept males and females, precluding any interactions between the sexes. We thus wondered if the previously reported phenomenon of ‘harmonic convergence’ was an epiphenomenon of the observed daily variations of male/female flight tone ratios around a center value of 1.5 and not a signature of an acoustic interaction between male and female mosquitoes during paired flight (see also online supplement).

‘Harmonic convergence’ describes the transient (1-2s long) match of the third harmonic of the female flight tone with the second harmonic of the male flight tone. At a (male:female) fundamental flight tone ratio of 1.5, harmonic convergence will occur by default, independent of any interaction. We therefore conducted an in-depth analysis of the only existing, extensive, and publicly available, experimental data set (*22*) that had previously been generated to allow for a statistical perusal of harmonic convergence events in mosquitoes (see Figs. 4, S9-S16 and online supplement). We found that the number of harmonic convergence events between a specific male/female pair was only a function of their respective median flight tones, more specifically of the distance (or proximity) of the ratio of their median flight tones to a chosen harmonic convergence ratio (Tab S1), e.g. the ratio of 1.5. The closer a particular pair (virtual or real) was to the 1.5 ratio, the more harmonic convergence events occurred by mere chance (Fig. 4C and online supplement). Harmonic convergence events were not enriched in real pairs as compared to virtual pairs (composed of pairs chosen randomly from pools of lone flying males and females). Harmonic convergence events were also not more likely to occur in males exposed to playbacks of female flight tones (number of convergence events per minute, median±SE: real/live pairs= 3±0.56; virtual/loner pairs=4±0.78; playback=2±0.55).

**Fig. 4.**
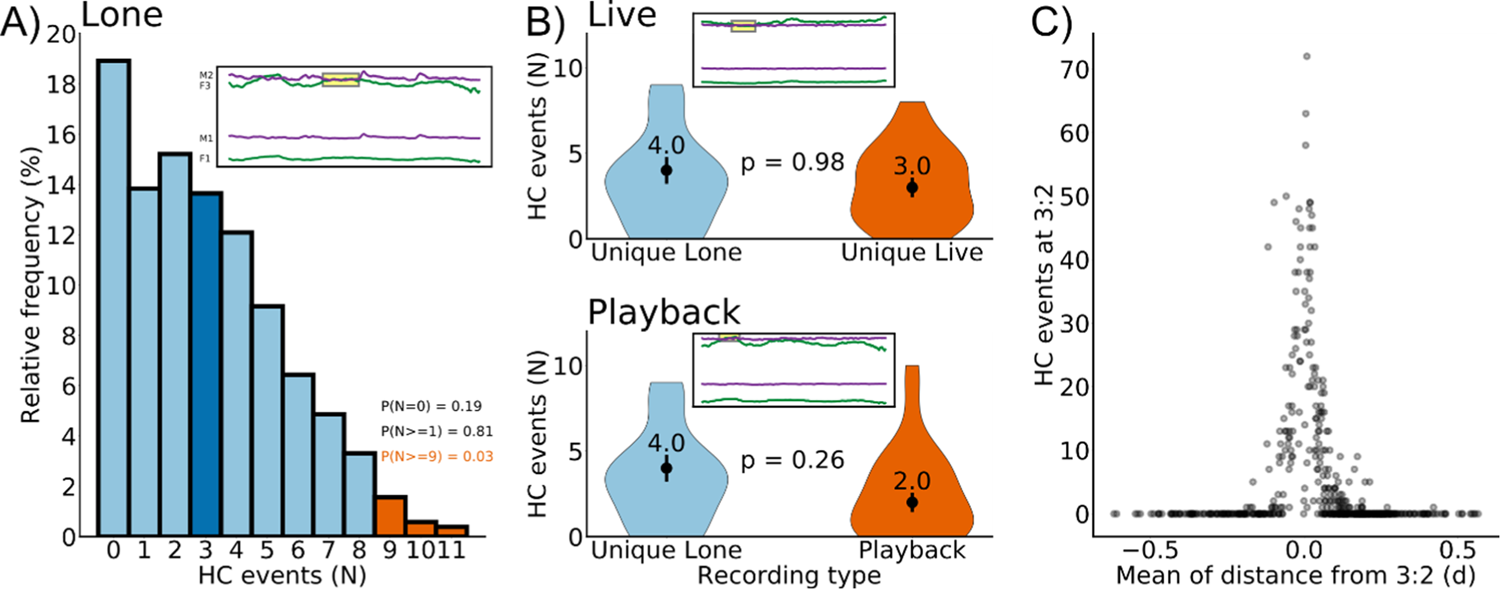
Statistical analysis of harmonic convergence (from data set A, Aldersley et al., 2016). Relative frequency distribution of the number of harmonic convergence events (N) counted for 513 virtual (‘lone’) pairs. This distribution serves as reference (null hypothesis) to test if harmonic convergence events are significantly different in real (‘live’) pairs. For an individual harmonic convergence event to be of statistical significance (i.e. to have a probability p<0.05 of having occurred by chance), a live pair must exhibit at least 9 harmonic convergence events during a one-minute long flight. Any value below 9 does by itself not constitute a statistically noteworthy event (i.e. it has a probability p>0.05 of having occurred by chance). The average number of harmonic convergence events (see also Fig. S14 and S15) for both lone and live pairs is ∼3. The inset illustrates an example of harmonic convergence at 3:2 for a lone pair. B) Violin plots of the number of harmonic convergence events observed in live pairs (top) and playback pairs (bottom) compared to the number of harmonic convergence events in unique lone pairs. No statistical difference is observed between the groups. Insets illustrates an example of harmonic convergence of a live pair (top) and a playback pair (bottom). C) An illustration of how the number of harmonic convergence (HC) events at a given ratio exhibited by a mosquito pair is a function of the pair’s mean distance (*d*) from that ratio, whereby *d* is calculated as mean of the absolute distances of a pair’s instantaneous flight tone ratios (across the one minute flight) from the given harmonic ratio *Hr* (here 3:2). Note the sharpness of the distribution peak, which indicates an extreme noise sensitivity of HC calculations from pairs whose flight tone ratios are accidentally close to the HC ratio of interest (here 3:2); such pairs will produce large numbers of HC events by mere chance. In contrast, pairs whose flight tones are accidentally far from the HC ratio will not produce any convergence events, at all.

We also tested if flight tones of lone-flying mosquitoes (males and females) were in any other way different from those of flying in pairs (Table S2 and S3). But we found no significant differences in flight tone frequency or variance; all tested cohorts were statistically indistinguishable from each other. In summary, there is no evidence for acoustic interaction between the sexes; all occurring harmonic convergence events were sufficiently explained by chance. This was true for both males and females and for all other harmonic convergence ratios suggested (on this point see also below).

### Mosquito flight tone *phonotypes*

We tested if the swarming related - and male-specific - flight tone modulation seen in groups would also occur in mosquitoes kept individually. As observed under grouped conditions, the flight tones of individual females did not show significant differences at swarm time as compared to the rest of the day (Fig. 5A; swarm time = 616.92±20.98 Hz, other time = 609.48±17.18 Hz; n = 18; p = 0.15). Individual males, in contrast, showed a significant increase at swarm time (Fig. 5A; swarm time = 908.94±31.91 Hz, other time = 886.75±29.37 Hz; n = 18;p = 0.031). When comparing the flight tone ranges of lone-flying mosquitoes, it was evident that both males and females occupied narrower frequency ranges than the population of all individuals of the respective sex pooled together (Figs. 5B, S6A). Within each sex, the flight tones of individual mosquitoes showed statistically significant differences (Kruskal-Wallis rank sum test, p<2.2e-16 in both males and females). This is remarkable - and relevant - from an acoustic perspective. The audibility of a given female for a given male depends on the extent of audible distortion products (DPs) that are generated by the mixing of his own flight tone with the flight tone of that particular female (Fig. 5C, left). For an ideally tuned male auditory nerve (see Figs. 3), the audibility score of a specific female can be approximated by calculating the overlap between quadratic and cubic DPs generated by the two flight tones.

**Fig. 5.**
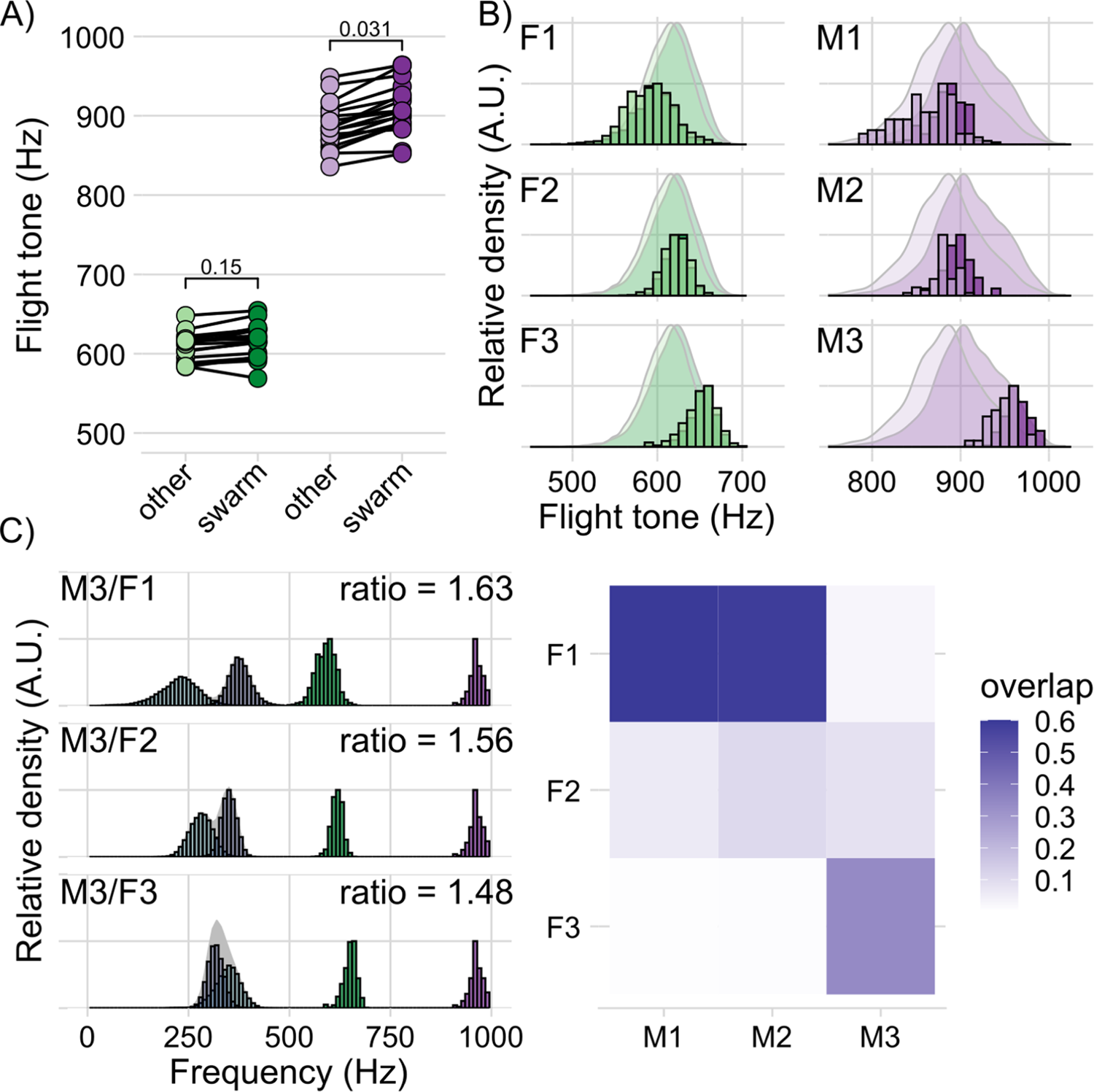
Single mosquito phonotype analysis. A) Swarm and out-of-swarm (other) flight tones of individual female (green) and male (purple) mosquitoes recorded during 12-hour light/dark entrainment (with 1 hour simulated dusk and dawn) at 28°C. Points are median flight tones calculated from all detected flight tones in the designated time window for each individual (F-other = 609.48 Hz, σ = 17.18 Hz, F-swarm = 616.92 Hz, σ = 20.98 Hz n = 18 females; M-other = 886.75 Hz, σ = 29.37 Hz, M-swarm = 908.94 Hz, σ = 31.91 Hz n = 18 males). B) Flight tone distributions of three representative female (F1-F3) and three representative male (M1-M3) phonotypes. Bar plots are scaled, binned counts of the individuals flight tones recorded during swarm time (darker color) and out-of-swarm time (lighter color). These are plotted against the population scaled density plots for comparison. C) Distribution of distortion tones (cubic – light blue, quadratic – dark blue) that would be produced between M3 (purple) and the three representative females (F1-F3, green). Average ratio value is calculated for every combination of male and female flight tone for each pair (M3/F1 = 1.63, σ = 0.07, n = 65,160 pairs; M3/F2 = 1.56, σ = 0.04, n = 33,240 pairs; M3/F3 = 1.48, σ = 0.04, n = 41,280 pairs). Heatmap displays the proportion of overlap between the two calculated distortion tone distributions for each representative female/male pair. Both female and male data are pooled from 3 independent experiments each with 6 individuals.

The observed individual differences between male and female flight tones - which partly form discrete, narrowband, and non-overlapping, *phonotypes* - are bound to lead to individual hearing ranges, which mean that some males can hear some females better than others (see Fig. 5C, right). For low sample sizes, the existence of narrowband *phonotypes* also introduces distinct peaks into the histograms of flight tone – or flight tone ratio – distributions (see supplement for details). This ‘peakiness’ of the landscape of male/female flight tone ratios has previously been interpreted as a signature of acoustic interactions and led to the postulation of additional ‘harmonic convergence’ ratios beyond 1.5 (*22*). Our analyses did not find any statistical evidence supporting such a conclusion, the suggested additional ratio peaks also disappear after averaging across appropriate sample sizes.

## Discussion

### Male control of female audibility

The acoustic chase of flying females, is a hallmark of reproductive behavior in male mosquitoes. Yet a common, and rather surprising, denominator of males across species, is that their auditory nerves are near-deaf to the actual flight tones of their conspecific females (*25*) (Figs. 3B, S5).

Females will become audible to males, however, if the nonlinear mixing of male and female flight tones produces audible distortions within the male’s ear. Here, the degree of female audibility depends on the specific interrelation of the two flight tones. We found significant daytime- and state-dependent modulations of flight tones in single-caged males, but not in single-caged females. The respective audibility space of the mating swarm is under circadian modulation and controlled by males. The detailed quantitative analysis of this audibility control shows how the males’ flight tone (or wing beat frequency) selection exploits the boundaries of their hearing ranges (Fig. 3A,B and S5), revealing a close coupling between the ranges of mosquito wing beat frequencies and the frequency response functions of their flagellar ears.

Theoretical considerations predict an optimality of distortions to occur around a (male:female) frequency ratio of 1.5. At this optimal ratio, the number of produced distortions would peak at frequencies equal to half the female’s flight tone (*f*_1_/2). In fact, simply dividing the recorded female flight tones by 2 would provide a good estimate for the center frequency of male auditory nerve responses (see dotted line in Fig. 3). Multiplying the response plateau of the male nerve by 2, in turn, can predict the distribution width of female flight tones. Notably, while *Anopheles* males optimize their flight tones for female detection in the *swarming* state, a residual audibility of females is maintained in the *baseline* state (Fig. 3). The here relevant evolutionary pressures on flight tone selection merit further exploration. The wing beat frequency upshift that associates with the *activated* state (i.e. after hearing a female) seems to move the males away from the optimal ratio to female flight tones. This could indicate that during the copulatory chase males sacrifice female audibility to flight speed. An alternative explanation might be that males shift their flight tones in order to be less audible to the female, thus flying in ‘stealth mode’. It could, finally, be that – during the actual chase - females show a wing beat frequency upshift themselves (preliminary data from Fig. S8 is consistent with this hypothesis) – maybe also accompanied by an increase in flight speed; this might restore the optimal 1.5 ratio and render the females more audible again (see Fig. S8) - but also more difficult to catch. Whatever the ultimate reason, the observed flight tone changes are bound to affect the mutual audibility in a hearing system, where tonal changes of ∼50Hz can reduce responses by >60% (*28*).

These general settings hold true for both *Anopheles gambiae* and *Aedes aegypti* (Fig. S4), but distinct differences exist in their behavioral, particularly their circadian, activity patterns (compare Figs. 2 and S4) (*29*). Males of *Anopheles* show a narrow pattern of activity (and phonotactic responsiveness) (Figs. 1 and 2), almost exclusively occurring at swarm time (ZT13, sunset). *Aedes* males, in contrast, display a much wider range of daily activity, although interestingly peaks in activity remain at dawn and dusk (*30*) (Fig. S4). In alignment with these behavioral patterns, the flight-tone-mediated sensitization to female flight tones is restricted to ZT13 in *Anopheles*, whereas males of *Aedes* appear to remain optimally sensitive to female sounds for larger parts of the day (Fig. S7).

Flying with higher flight tones – and thus at higher wing beat frequencies – can be expected to be costly. Any increase in female audibility that males can gain from beating their wings faster must thus be traded off against the corresponding energetic costs. We assume that the differences seen between the two species reflect differential reproductive strategies. *Anopheles gambiae* males mate mostly in large (hundred- to thousand-strong) swarms, which only form once a day at dusk. *Aedes* males mate more widely across the day and form only small swarms (with dozens of individuals), which also form at dusk (*29*).

Rather than reflecting acoustic interactions - or communication - between males and females, mosquito flight tone variations are linked to male-specific adjustments of female audibility. Depending on their circadian or behavioral state, males retune the ‘pitch’ of produced distortions, thus modulating female audibility and increasing (or decreasing) the likelihood of reproductive interactions.

Particularly intriguing in this context are the possible trade-offs between flying and hearing. It has been reported that the upshift in male wing beat frequency after playback of female flight tones coincides with an increase in flight speed (*27, 31*). The males’ phonotactic search thus receives both aerodynamic and acoustic support. The underlying biophysics of these relations are non-trivial. Forcing a simple harmonic oscillator beyond its best frequency leads to a reduction in oscillation amplitude (e.g. here the wing stroke angle) and thus – everything else equal - a reduction in flight speed. Mosquitoes, however, appear to have evolved a different mechanism of force generation, related to wing rotation (*32*) which enables higher wing beat frequencies. It is tempting to speculate that this unique mode of operation evolved to partly uncouple the aerodynamic and acoustic roles of mosquito wing beats.

The existence of individual (male and female) phonotypes, finally, suggests that female audibility differs across males, with some females being a better ‘acoustic match’ for a given male than others. The population-genetic consequences of such inter-individual variance in mating compatibility could be substantial. While the molecular bases for mosquito flight tone frequencies are still largely unknown, recent work has linked first genetic pathways and also demonstrated the behavioral impact of mutant phenotypes (*28*).

The acoustic fitness of males forms a crucial bottleneck for their reproductive success; it’s correct assessment requires the knowledge, and triangulation, of the spectro-temporal properties of three elements: (i) the males’ own flight tones, (ii) the females’ flight tones and (iii) the response function of the males’ flagellar ears (both mechanically and neuronally). On the level of individual pairs, this will allow for a quantitative understanding of mate selection (e.g. female choice/male choice). On the population level, this will not only contribute insights into mosquito evolution but also help optimize mosquito mutants for mass release programs (e.g. *gene drive*). ‘Harmonic convergence’ events, finally, emerge as epiphenomena from the rules that govern the mosquitoes’ distortion-centered flight tone variance. Future mosquito research will need to keep at least one eye on distortions to see clear.

## Acknowledgments

We thank Alexandros Alampounti for valuable discussions and advice.

## Author contributions

Conceptualization, J.S., M.G., M.P.S., and J.T.A.; Investigation, J.S., M.P.S., M.A., W.N., J.B.; Writing—Original Draft, J.S., and J.T.A.; Methodology, J.S., M.G., M.P.S., and J.T.A.; Writing—Review & Editing, J.S., M.P.S., C-H.C. and J.T.A.; Visualization, J.S., M.G., M.P.S., and J.T.A.; Supervision, J.T.A., J.S.; Software, J.S., M.G., M.P.S.; Resources, J.S., G.M., S.M., J.B., R.S., M.P.S., C-H.C. Funding Acquisition, J.T.A., S.M., M.A.;

## Data and materials availability

All data is available in the main text, the supplementary materials or can be made available upon request.

## Supplementary Materials

### Materials and Methods

Annex 1: Statistical analysis of harmonic convergence events (data from *Aldersley et al*. 2016) including Figures S9-S16

### Materials and Methods Insect rearing

Mosquitoes were reared at 28°C, 80% following standard protocols. Briefly, eggs laid onto filter paper were floated using 1% w/v tonic salt solution (Blagdon Pond Guardian Tonic Salt). Larvae were feed on increasing amounts of TetraMin fish flakes (Tetra®) and were split to maintain density with water being replenished as required. Pupae collected daily, were sexed into 50mL falcon tubes, and following eclosion, the adults were then transferred to experimental arenas.

### Activity monitoring

One (single-housed) or three (group-housed) mosquitoes were aspirated into glass activity tubes (125mm long, 25mm ⌀) covered at both ends with cotton wool, with one end soaked in 10% sucrose solution and wrapped in parafilm to maintain a food source and humidity. Activity tubes were then loaded in TriKinetics LAM25 locomotion activity monitors inside of a Percival I- 30VL environmental chamber. Activity counts were recorded as infrared beam breaks as the mosquito(es) crossed the midline of the tube, counts were binned at one-minute intervals.

Double plotted actograms and activity plots were created using the Rethomics R package (*33*). Free running period was calculated using chi^2^ periodogram analysis to create a sliding window for peak activity analysis.

### Flight tone recordings

Approximately 100 virgin individuals were aspirated into a BugDorm-1 insect rearing cage that was then placed in a Percival I-30VL environmental chamber. Chamber conditions were set to a constant temperature of 28°C, 80% relative humidity and a 12-hour light/dark cycle with a 1- hour ramped light transition period to simulate both dawn and dusk (the onset of each of these transition periods will be referred to as ZT00 and ZT12 respectively). Audio recordings were then taken using the microphone array placed in the middle of the BugDorm, recordings were either performed at specific times of the day or continuously depending on the experiment.

### Data acquisition

For swarm recordings, four particle velocity microphones (Knowles NR-23158-000) were arranged in a cube in order to capture mosquito flight tones from all directions. The cube was attached to a 4mm ⌀ rod that was fixed in the centre of a 300mm^3^ BugDorm-1 insect rearing cage. For individual recordings, a single microphone was fixed in the centre of a 3D printed 50mm^3^ cage. Microphones were then connected to custom made preamplifiers (Uni Köln - DETAILS) which were then connected to a CED 1401 (Cambridge Electronic Design) data acquisition interface. Experiments involving artificial tone playback, had a small speaker driver near the microphone which was triggered using the DAC output channels of the CED 1401 device. Audio was recorded at 50kHz per channel and flyby events were identified and quantified using an automated pipeline.

### Signal Processing

Recorded data was segmented into 1-minute chunks before processing. Any DC component was removed before passing the signal through a digital bandpass filter (4^th^ order, Butterworth design). Corner frequencies were 300 & 1200Hz except in experiments where artificial female flight tones were played in which corner frequencies were 600 & 1200Hz. A moving average envelope was computed across the fully rectified signal in order to identify flyby events (i.e. the envelope is >2*rectified local mean). Identified flyby events were then fitted with a sliding general sine wave model (10ms window, 50% slide) to extract frequency information. Medians of all fits that had a R^2^ > 0.9 were calculated to quantify the main frequency component of each flyby event.

Despite signal processing efforts, in some experiments, mechanical background noise from the environmental chamber were detected by the pipeline, often observed as regular, short events (∼30ms) across a very narrow frequency band (μ = 653Hz & σ = 3.7Hz). Minimum event length was adjusted until no events were detected during the light phase of the experiment i.e. when Anopheles mosquitoes do not exhibit any flight activity. This value was typically set at > 30ms.

### Nerve recording methodology

4-8 day old mosquitoes were first sedated on ice and mounted on Teflon rods using blue-light cured dental glue. After application of this glue, only the right flagellum was free to move. The rod was then held in place by a micromanipulator on top of a vibration isolation table, with the mosquito orientated to face a laser Doppler vibrometer at a 90° angle.

In order to provide electrostatic stimulation to the mosquito ear, a charging electrode was inserted into the mosquito and its electrostatic potential was increased to −20 V relative to the ground. Two electrostatic actuators were placed symmetrically around the flagellum to enable ‘push and pull’ electrostatic stimulation. A recording electrode was then inserted at the base of the right pedicel so recordings could be made of compound antennal nerve responses to stimulation. Flagellar displacements resulting from stimulation were simultaneously recorded with electrophysiological activity using the vibrometer. At both the beginning and end of the stimulation, free fluctuation recordings were taken of the flagellum to test for changes in auditory capabilities during the experiment.

Stimulation came in the form of mono-frequent pure tones (sine waves) played sequentially from 15 to 695Hz in 10Hz intervals. Each stimulus playback lasted 2.5 seconds, followed by a further 2.5 seconds of silence before the next stimulus began. Each stimulus frequency was played five times.

Analysis of nerve responses to pure-tone stimulation is complicated because of potential crosstalk between the stimulus and the recording electrode set up, which can result in reflections of the stimulus amplitude in the nerve response. These artifacts can artificially distort apparent nerve responses. One way of counteracting this is to take advantage of the known phase of the artifact, as well as its’ distinguishability from the real nerve response because of the lack of a time delay. We assumed that if the crosstalk present in the system results in an overlaying of a copy of stimulus into the nerve response, then by subtracting this artifact (after accounting for a change of scale) only the real data should remain.

Stimulus and nerve data were therefore analysed using a custom MATLAB script. A DC remove was applied to the steady state responses of both sets of data, before each stimulus was subtracted from the corresponding nerve response and the remaining area under the curve analysed. To account for the change in scale between the two channels, the stimulus data was multiplied by a constant which was selected such that the area under the curve of the remaining nerve response was minimised. The nerve response in the absence of crosstalk could then be estimated from the power spectrum of the residual nerve response. The median best frequency of the nerve was defined as the frequency at which the nerve response magnitude was greatest.

In total, 8 Ae. aegypti females, 10 Ae. aegypti males, 7 An. gambiae females, 7 An. gambiae males, 8 Cx. quinquefasciatus females and 8 Cx. quinquefasciatus males were included in the final analysis.

### Data analysis

Activity and flight tone data analysis was performed using R 4.0.3 using a number of published packages from CRAN and custom scripts. Signal processing pipeline available from https://github.com/jaspwn/simbaR. Electrophysiology data analysis was performed using MATLAB using packages and custom scripts.

**Fig. S1.**
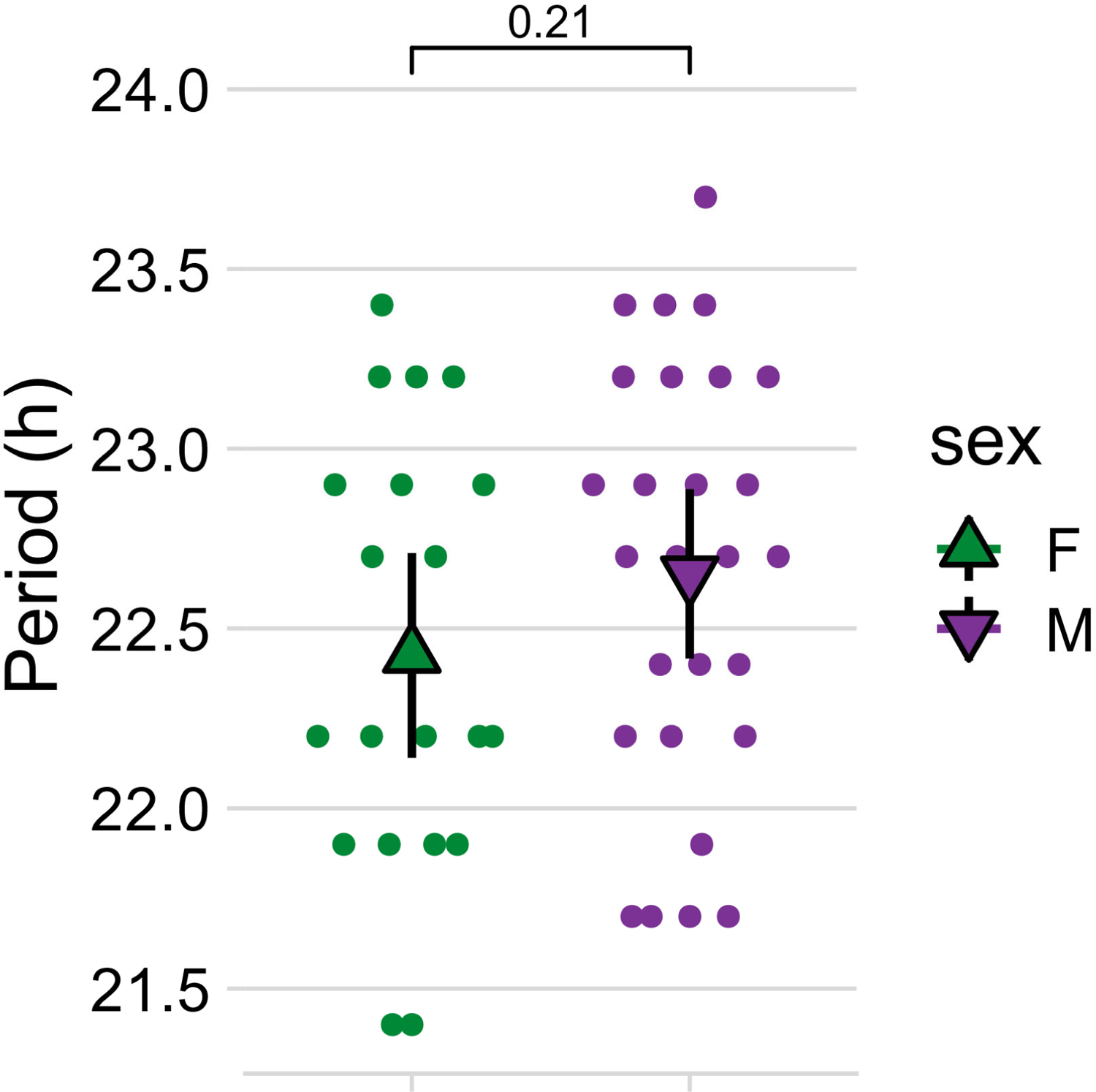
Period calculations of circadian locomotor pattern. Chi-squared periodogram analysis was used to estimate period length for each group during the free-running phase of the experiment (constant conditions). Points show individual calculated periods and triangles show the group mean for each sex ± S.E.M. (females = 22.43 hours ± 0.14 hours, n = 20; males = 22.65 hours ± 0.11 hours, n = 27). No significant difference in period is observed (*p* = 0.21, Welch Two Sample t-test).

**Fig. S2.**
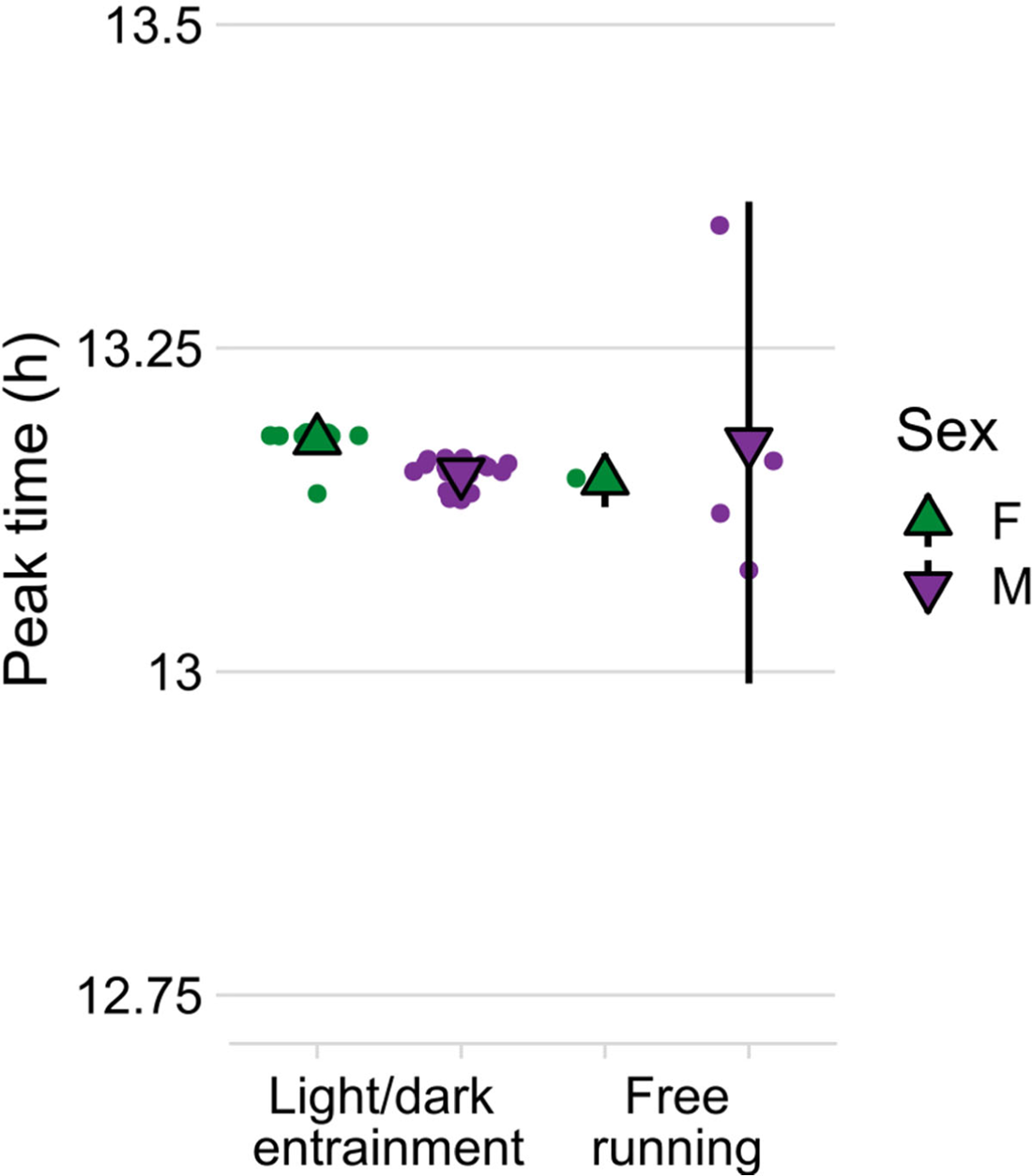
Daily peak activity of males and females quantified by the maximum number of flyby events past a stationary microphone. Points show daily peak activity and triangles show the group mean for each phase-sex combination ± S.E.M. LD-female = 13.18 h ± 0.004 h S.E.M., n = 12 days; LD-male = 13.15 h ± 0.002 h S.E.M., n = 18 days; FR-female = 13.15 h ± 0.001 h S.E.M., n = 2 days; FR-male = 13.29 h ± 0.05 h S.E.M., n = 4 days.

**Fig. S3.**
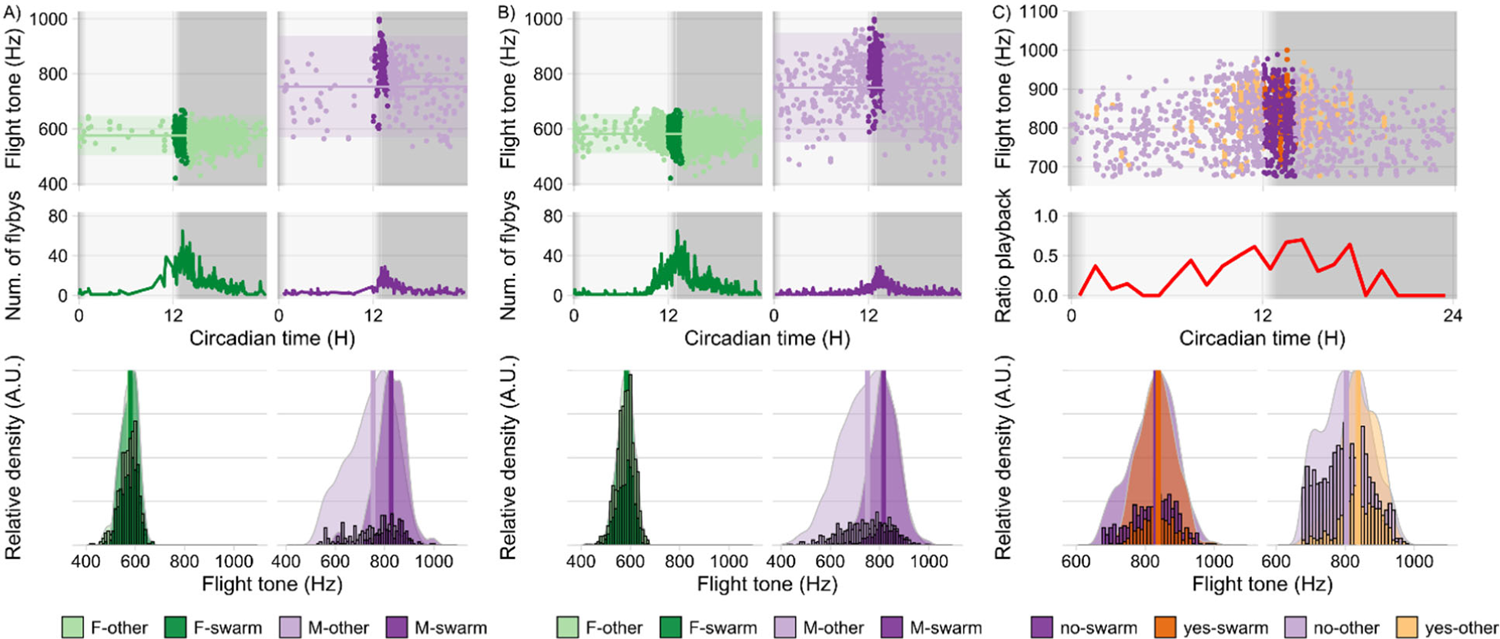
Acoustic analysis of free-flying populations of male and female *Anopheles* mosquitoes in free-running conditions A) *Top panels* Individual flight tones recorded from free-flying populations of females and males during the first day of constant darkness at 28°C. Points represent median flight tones of an individual flyby events (see Materials & Methods), darker colors indicate the swarm time period. Solid horizontal line shows the population median for out-of-swarm flight tones ± 95% C.I. for each sex (F = 576.69 Hz, σ = 35.59 Hz, n = 989 flight tones; M = 793.92 Hz, σ = 90.77 Hz, n = 395 flight tones). *Middle panels* Line plots showing running averages (window: 5 min) of number of recorded flight tones across the day as a measure of flight activity. *Bottom panels* Distribution of flight tones recorded for each sex across the entrainment days separated by phase – swarm & out-of-swarm (other). Bar plots are binned counts of the individual flight tones and they are plotted against scaled density plots in order to visualise the distribution shift between the swarm and out-of-swarm (other) group. Vertical lines indicate the calculated median for each sex-phase combination (F-swarm = 580.04 Hz, σ = 35.37 Hz, n = 424 flight tones; F-other = 575.61 Hz, σ = 35.64 Hz, n = 565 flight tones; M-swarm = 826.72 Hz, σ = 63.97 Hz, n = 162 flight tones; M-other = 753.52 Hz, σ = 92.30 Hz, n = 233 flight tones). B) *Top panels* Individual flight tones recorded from free-flying populations of females and males over several days in constant darkness at 28°C. Points represent median flight tones of individual flyby events (see Materials & Methods), darker colors indicate the swarm time period. Solid horizontal line shows the population median for out-of-swarm flight tones ± 95% C.I. for each sex (F = 582.38 Hz, σ = 35.08 Hz, n = 2347 flight tones; M = 780.39 Hz, σ = 96.64 Hz, n = 1011 flight tones). *Middle panels* Line plots showing running averages (window: 5 min) of number of recorded flight tones across the day as a measure of flight activity. *Bottom panels* Distribution of flight tones recorded for each sex across the entrainment days separated by phase – swarm & out-of-swarm (other). Bar plots are binned counts of the individual flight tones and they are plotted against scaled density plots in order to visualise the distribution shift between the swarm and out-of-swarm (other) group. Vertical lines indicate the calculated median for each sex-phase combination (F-swarm = 583.69 Hz, σ = 34.18 Hz, n = 762 flight tones; F-other = 581.49 Hz, σ = 35.5 Hz, n = 1585 flight tones; M-swarm = 816.44 Hz, σ = 66.80 Hz, n = 300 flight tones; M- other = 750.00 Hz, σ = 99.30 Hz, n = 711 flight tones). C) *Top panels* Individual flight tones recorded from populations of free-flying male mosquitoes presented with a 1-minute artificial female flight tone (550Hz) at 30-minute intervals. *Middle panels* Line plot shows ratio between flight tones recorded during playback of artificial female flight tone and the total number of flight tones within each 30-minute interval (n_(playback)_/n_(total)_). *Bottom panels* Bar plots are binned counts of individual flight tones; they are plotted against scaled density plots in order to visualize the distribution shift between the playback and no playback group for each experimental phase. Vertical lines indicate the calculated median for each playback-phase combination (no-swarm = 830.20 Hz, σ = 64.98 Hz, n = 461 flight tones; yes-swarm = 837.56 Hz, σ = 51.00 Hz, n = 223 flight tones; no-other = 803.35 Hz, σ = 69.19 Hz, n = 1136 flight tones; yes-other = 838.24 Hz, σ = 56.32 Hz, n = 383 flight tones). Female data are pooled from 3 independent experiments, male data are pooled from 4 independent experiments and playback data are pooled from 4 independent experiments.

**Fig. S4.**
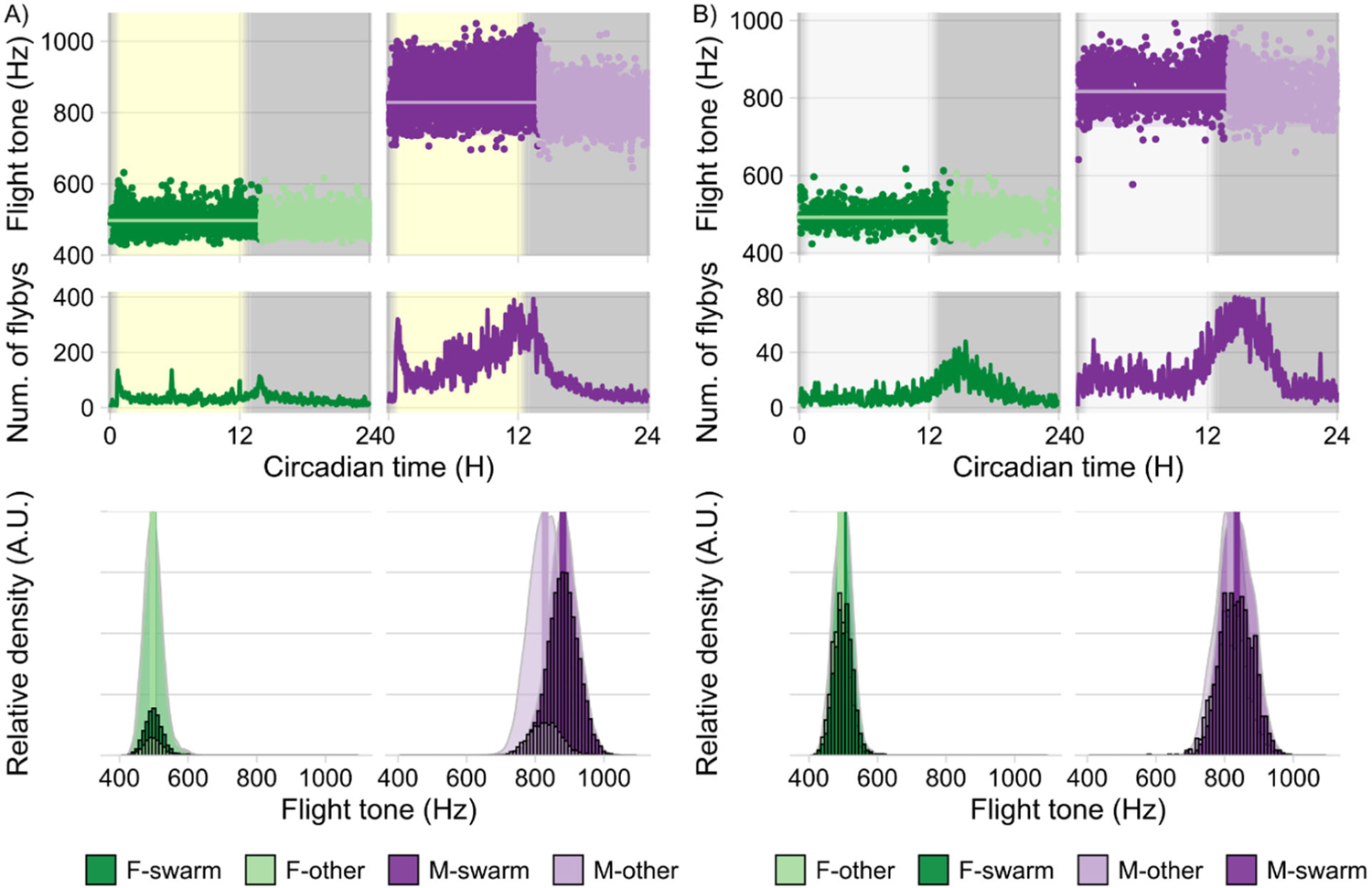
Acoustic analysis of free-flying populations of male and female *Aedes aegypti* mosquitoes. A) *Top panels* Individual flight tones recorded from free-flying populations of females and males during 12-hour light/dark entrainment (with 1 hour simulated dusk and dawn) at 28°C. Points represent median flight tones of an individual flyby events (see Materials & Methods), darker colors indicate the swarm time period. Solid horizontal line shows the population median for out-of-swarm flight tones ± 95% C.I. for each sex (F = 497.25 Hz, σ = 26.68 Hz, n = 952 flight tones; M = 828.88 Hz, σ = 46.95 Hz, n = 3202 flight tones). *Middle panels* Line plots showing running averages (window: 5 min) of number of recorded flight tones across the day as a measure of flight activity. *Bottom panels* Distribution of flight tones recorded for each sex across the entrainment days separated by phase – swarm & out-of-swarm (other). Bar plots are binned counts of the individual flight tones and they are plotted against scaled density plots in order to visualise the distribution shift between the swarm and out-of-swarm (other) group. Vertical lines indicate the calculated median for each sex-phase combination (F-swarm = 497.25 Hz, σ = 27.11 Hz, n = 2374 flight tones; F-other = 497.25 Hz, σ = 26.68 Hz, n = 952 flight tones; M-swarm = 880.95 Hz, σ = 45.02 Hz, n = 16220 flight tones; M-other = 828.88 Hz, σ = 46.95 Hz, n = 3202 flight tones). B) *Top panels* Individual flight tones recorded from free-flying populations of females and males in constant darkness at 28°C. Points represent median flight tones of an individual flyby events (see Materials & Methods), darker colors indicate the swarm time period. Solid horizontal line shows the population median for out-of-swarm flight tones ± 95% C.I. for each sex (F = 492.31 Hz, σ = 26.20 Hz, n = 784 flight tones; M = 816.77 Hz, σ = 46.05 Hz, n = 1252 flight tones). *Middle panels* Line plots showing running averages (window: 5 min) of number of recorded flight tones across the day as a measure of flight activity. *Bottom panels* Distribution of flight tones recorded for each sex across the entrainment days separated by phase – swarm & out-of-swarm (other). Bar plots are binned counts of the individual flight tones and they are plotted against scaled density plots in order to visualize the distribution shift between the swarm and out-of-swarm (other) group. Vertical lines indicate the calculated median for each sex-phase combination (F-swarm = 500.43 Hz, σ = 27.01 Hz, n = 808 flight tones; F-other = 492.31 Hz, σ = 26.20 Hz, n = 784 flight tones; M-swarm = 834.59 Hz, σ = 46.73 Hz, n = 1664 flight tones; M-other = 816.77 Hz, σ = 46.05 Hz, n = 1252 flight tones). Both female and male data are pooled from 2 independent experiments.

**Fig. S5.**
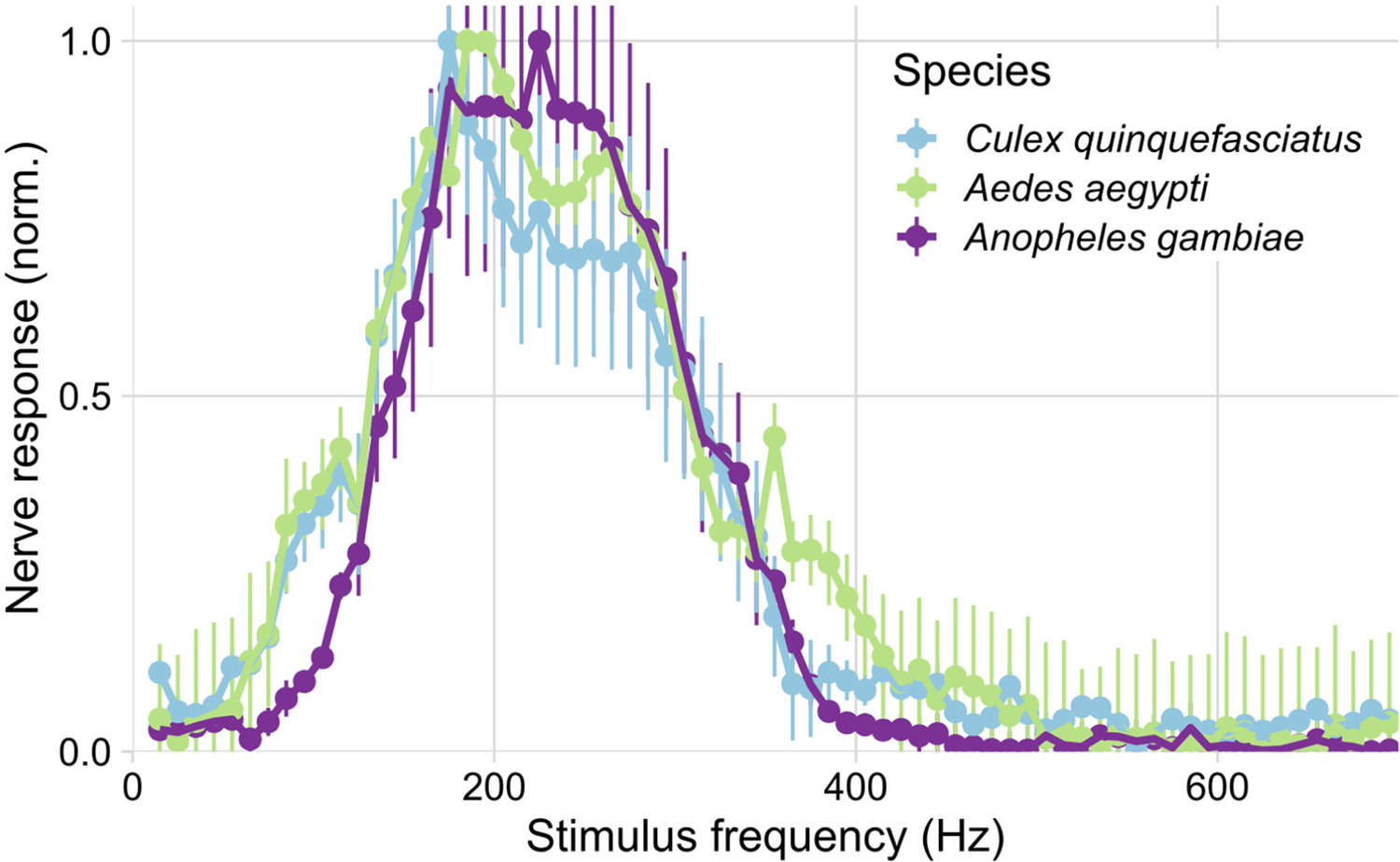
Normalized antennal nerve responses from three mosquito species to (electrostatic) pure tones played sequentially from 15 to 695Hz in 10Hz steps. Points are group medians from male antennal nerves for each species ± S.E.M (n = 8 *Culex quinquefasciatus* males; 10 *Aedes aegypti* males; 7 *Anopheles gambiae* males).

**Fig. S6.**
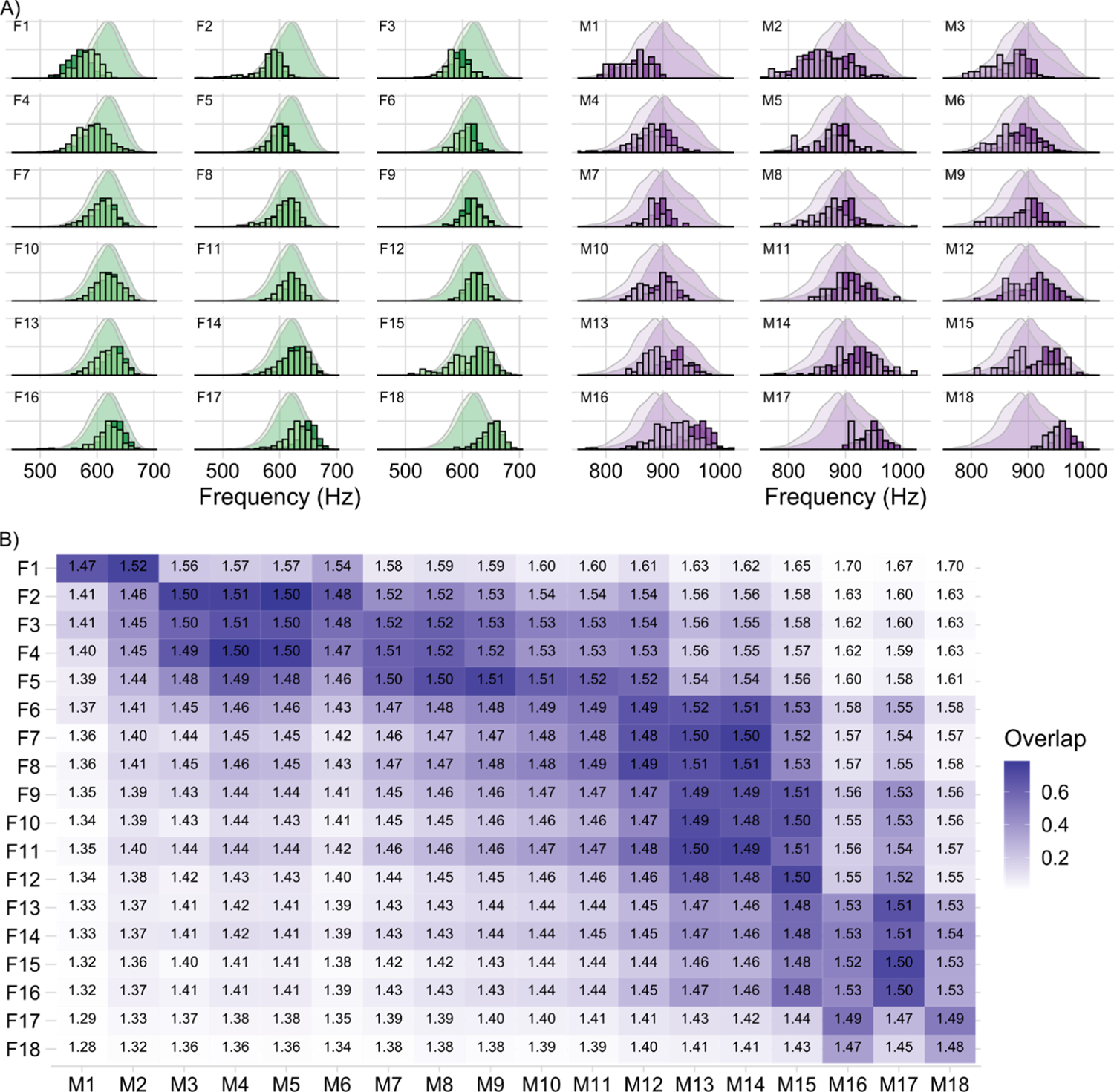
A) Flight tone distributions of all female (F1-F18) and all male (M1-M18) phonotypes. Bar plots are scaled, binned counts of the individuals flight tones recorded during swarm time (darker color) and out-of-swarm time (lighter color). These are plotted against the population scaled density plots for comparison. B) Heatmap displays the proportion of overlap between the two calculated distortion tone distributions for each female/male pair. Inset number is the average ratio value calculated for every combination of female and male flight tone for each pair Both female and male data are pooled from 3 independent experiments each with 6 individuals.

**Fig. S7.**
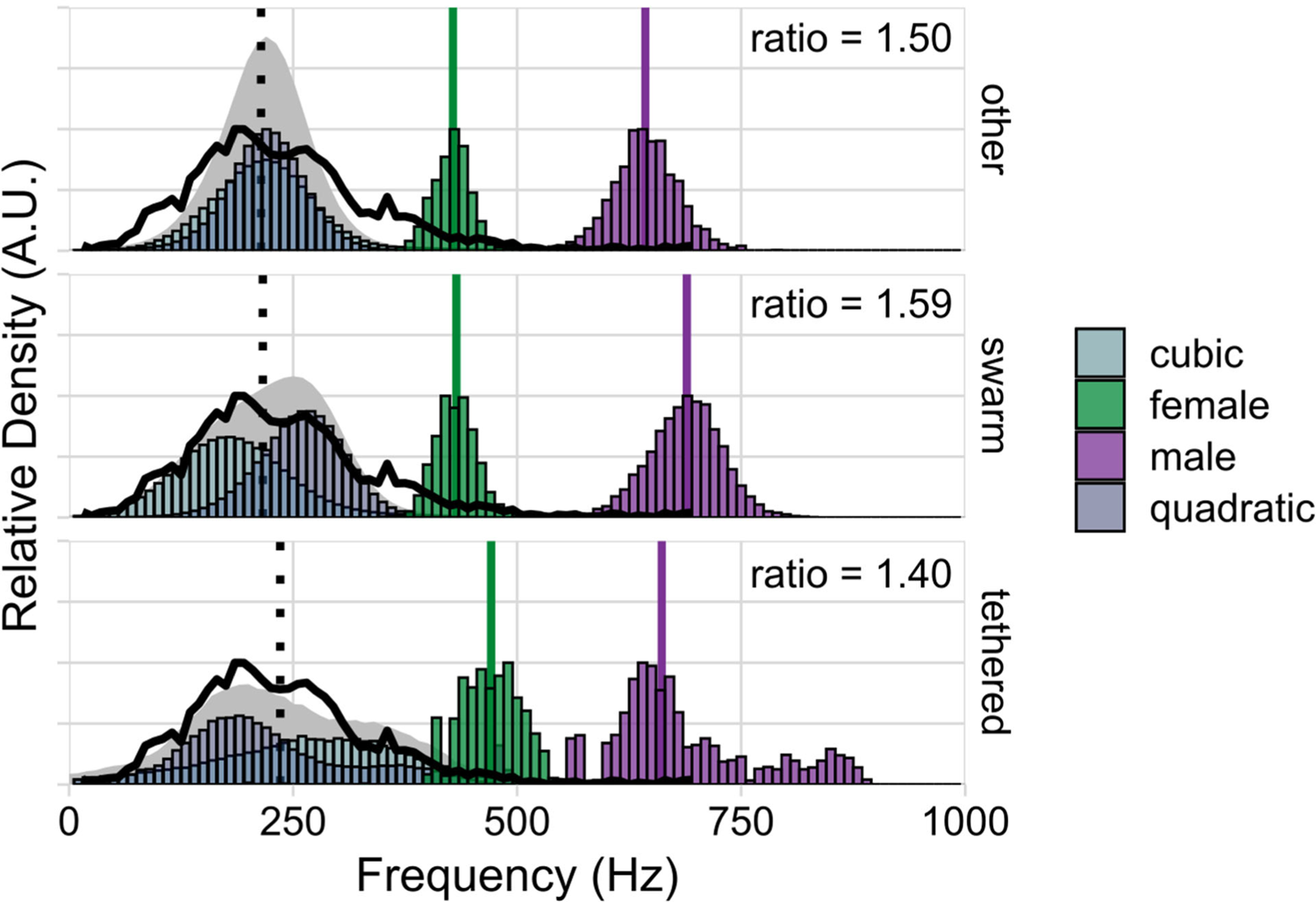
Predicted distortion products calculated for *Aedes aegypti* swarm cages at 22°C and tethered flight tone data we independently calculated from the data set A, *Aldersley et al., 2016*. At 22°C, we could match our own nerve recordings (solid black lines) to the created distortions. The vertical, dotted black line demarcates an optimum prediction for the center of male nerve responses for a distortion product based hearing system (optimum frequency = 0.5*f1). Note that - in contrast to *Anopheles* (see Fig. 3B) - *Aedes* remains at the optimal (male:female) flight tone ratio of 1.5 for most parts of the day (other); at swarm time, the male-specific wing beat frequency increase lifts the ratio to 1.59 (reminiscent of the ‘activated’ state in *Anopheles*); during tethered flight, in turn, the average ratio drops to 1.4. The colored vertical lines indicate the calculated median flight tone for each sex (other: females = 428.26 Hz, σ = 28.11 Hz, n = 670 flight tones; males = 643.11 Hz, σ = 36.13 Hz, n = 1644 flight tones; swarm: females = 432.22 Hz, σ = 27.23 Hz, n = 2243 flight tones; males = 689.35 Hz, σ = 40.85 Hz, n = 9704 flight tones; tethered: females = 471.02 Hz, σ = 35.93 Hz, n = 2280 flight tones; males = 661.20 Hz, σ = 91.75 Hz, n = 3237 flight tones). Average ratio values are calculated between these medians. Both female and male data are pooled from 2 independent experiments.

**Fig. S8.**
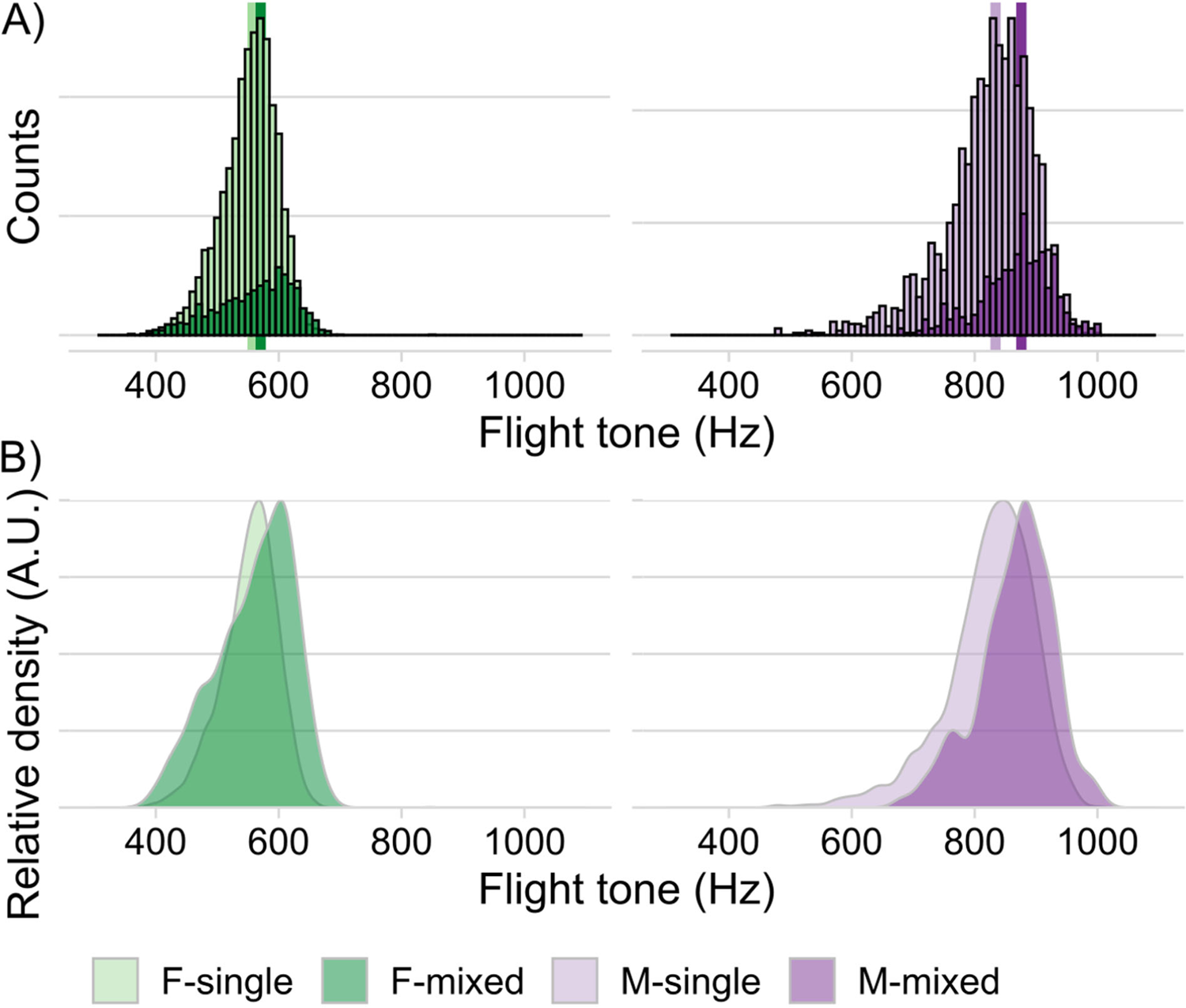
Comparison of single sex and mixed sex swarm cages of *Anopheles* mosquitoes (female, green; male, purple) recorded during 12-hour/12-hour light/dark entrainment (1-hour dusk/dawn transients) at 28°C. A) Bar plots are binned counts of individual flight tones scaled for visibility. A change in distribution can be observed for both sexes between single sex and mixed cages. Females appear to skew to higher frequencies but also increase the low frequency the tail of the distribution. Male distributions shift to higher frequencies with no apparent change in shape. Note that due to the apparent female frequency upshift in the mixed-cages, the male:female flight tone ratio would remain close to the optimum of 1.5. Vertical lines indicate the calculated median for each sex-condition combination (F-single = 557.33 Hz, σ = 45.36 Hz, n = 5379 flight tones; F-mixed = 570.45 Hz, σ = 63.13 Hz, n = 1422 flight tones; M-single = 834.05 Hz, σ = 72.67 Hz, n = 2044 flight tones; M-mixed = 876.07 Hz, σ = 61.04 Hz, n = 522 flight tones). Female single sex data are pooled from 3 independent experiments, male single sex data are pooled from 4 independent experiments and mixed cage data are pooled from 2 independent experiments. B) Conversion of flight tone counts to relative density plots to facilitate appreciation of distribution shifts.

**Fig. S9.**
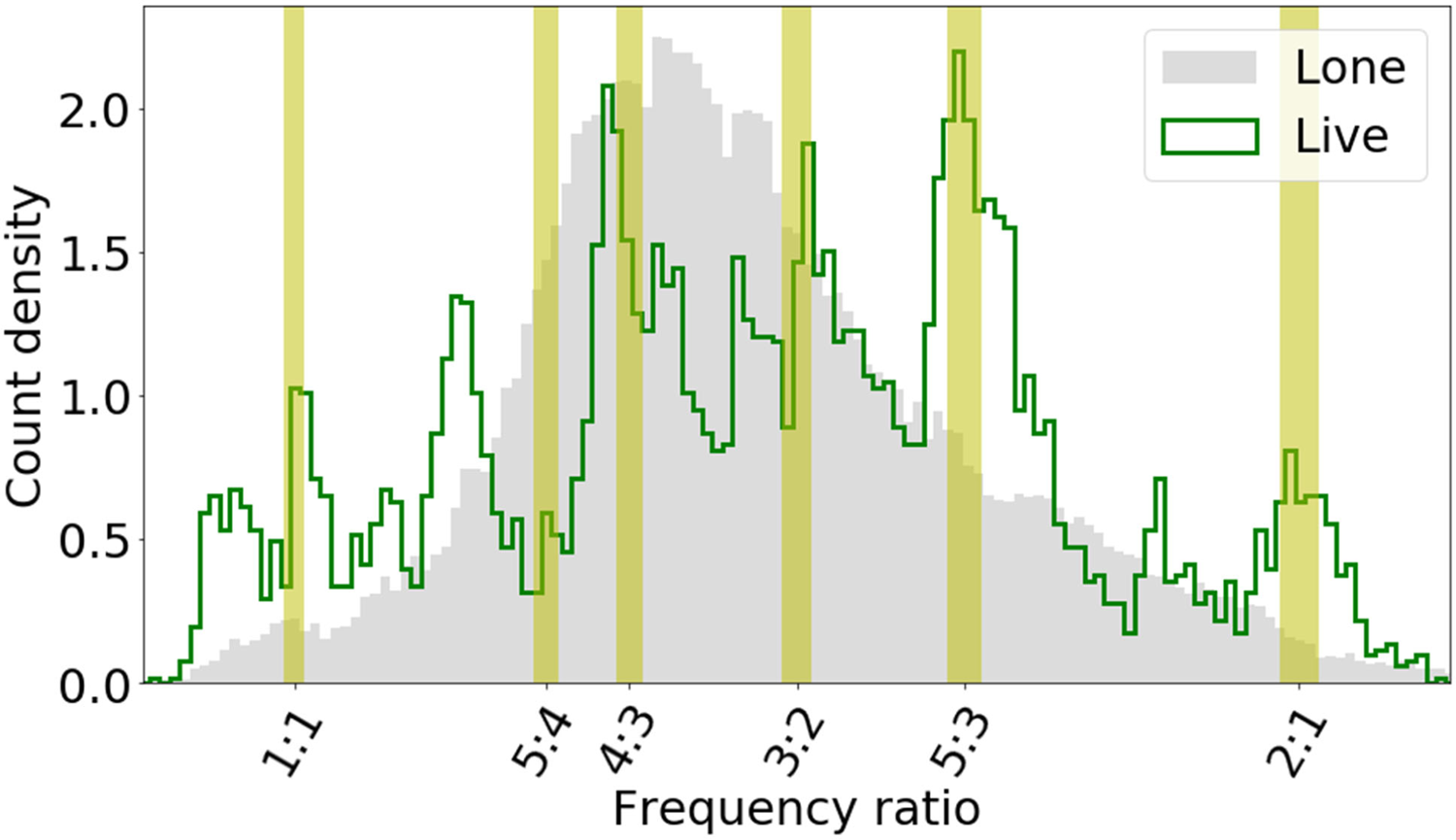
In a first step, we reconstructed the (male:female) flight tone frequency ratio histogram for tethered flights collected by *Aldersley et al., 2016* (*22*) for both live (real, green) and lone (virtual, grey) pairs of *Aedes aegypti* mosquitoes. Forty-three live and 513 lone pairs were used in total. The graph shows a smooth (∼unimodal) distribution for lone (virtual) pairs but a peaky (multimodal) distribution for the live (real) pairs, as also reported by *Aldersley et al., 2016* (*22*). *Aldersley et al., 2016* (*22*) compared the frequency ratio histograms of *live* and *lone* pairs, also with those of 34 live male-playback female pairs. These were constructed by stimulating 7 live males with subsets of 12 pre-recorded playbacks of females. We applied our analysis to the 34 pairs (here termed playback) to reconstruct the histogram that includes *live, lone* and *playback* frequency ratio distributions (bin width as above). Note that this is a part reconstruction of the original paper’s Figure panel 6b, as it does not include the frequency ratio distribution for live females stimulated with playbacks of males.

**Fig. S10.**
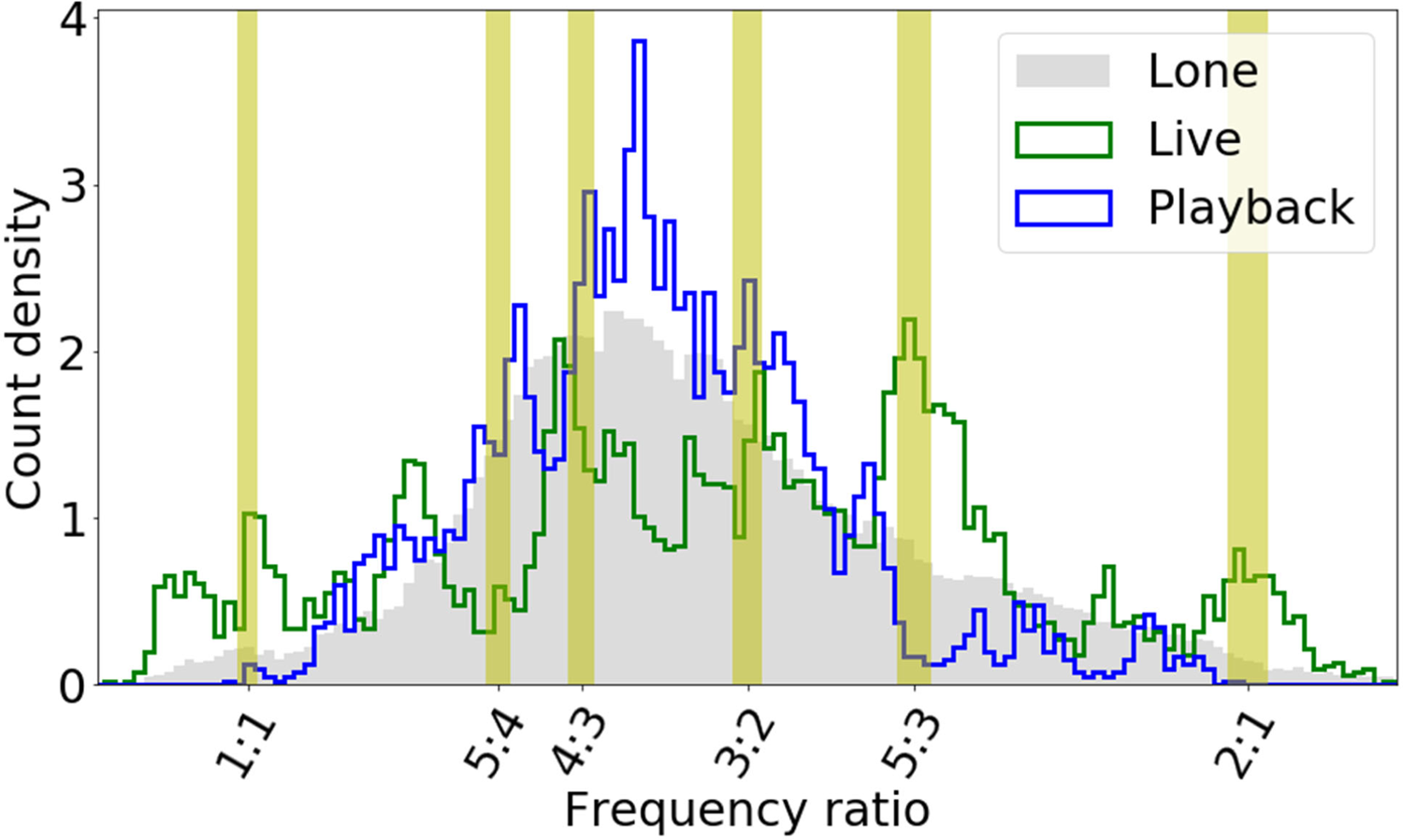
Reconstructed (male:female) flight tone frequency ratio histogram for tethered flights collected by *Aldersley et al., 2016* (*22*) for live (real, green), lone (virtual, grey), and pairs made of live males and virtual female stimulus (playback, blue). Forty-three live, 513 lone, and 34 playback pairs were used in total.

**Fig. S11.**
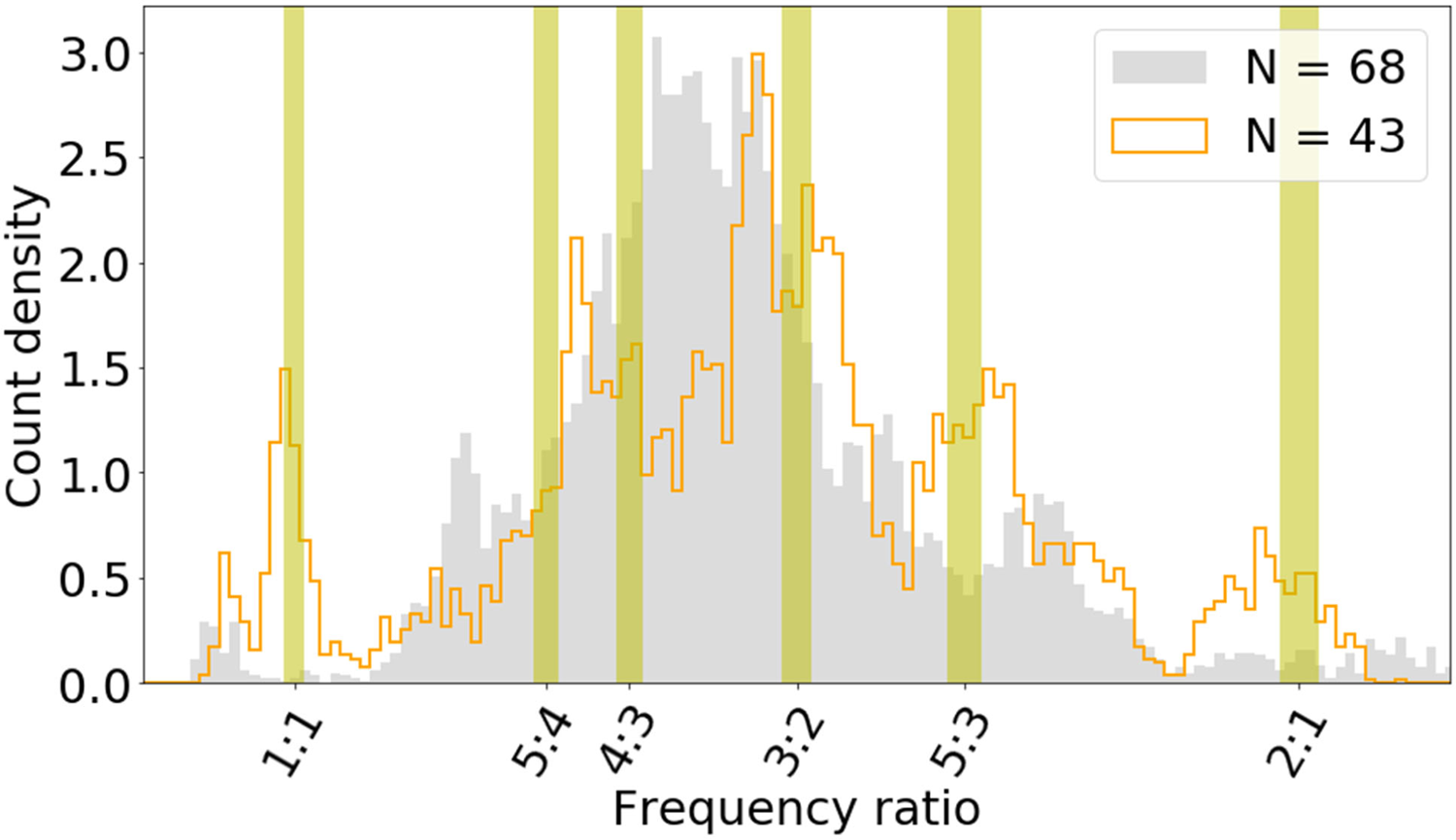
An illustration of how the number of pairs used can affect the shape of the distribution. These are qualitatively similar histograms to those of Figure S11, albeit produced using solely two subsets of lone pairs (with n= 68 and n=43, respectively) randomly selected from the total of 513. In an additional ‘bottom-up’ approach we analyzed the frequency - and frequency ratio - distributions of individual mosquitoes – and mosquito pairs from *Aldersley et al., 2016* (*22*) to trace the origins of the two different distribution shapes seen on the ‘population’ level. A look at the individual level shows that each mosquito (both males and females) occupies only a narrow range of flight tone frequencies (*Fig. S12, left*), much narrower than the flight tone distributions of all available samples combined would be. This phenomenon is equivalent to the ‘*phonotypes’* we observed for individual free-flying *Anopheles* (see Fig. 5 and S6). The narrow frequency ranges of individual males and females translate into sharp and narrow frequency ratios (*Fig. S12, right*), which in turn introduce sharp peaks into the population histograms. For low sample sizes (especially if re-using individual, or individual pairs of, mosquitoes) these distinct ratio peaks cannot be sufficiently averaged out and thus remain visible (as in *Fig. S9-S11* and Figure 6 of *Aldersley et al., 2016* (*22*)). The peaky spectral landscape of flight tone ratios reported for live pairs is thus introduced by the phonotypic nature of individual mosquitoes; it does not require, or reflect, any acoustic interaction between male and female. Rather, it is a phenotypic relic of a statistically incomplete, and thus non-representative, population sample.

**Fig. S12.**
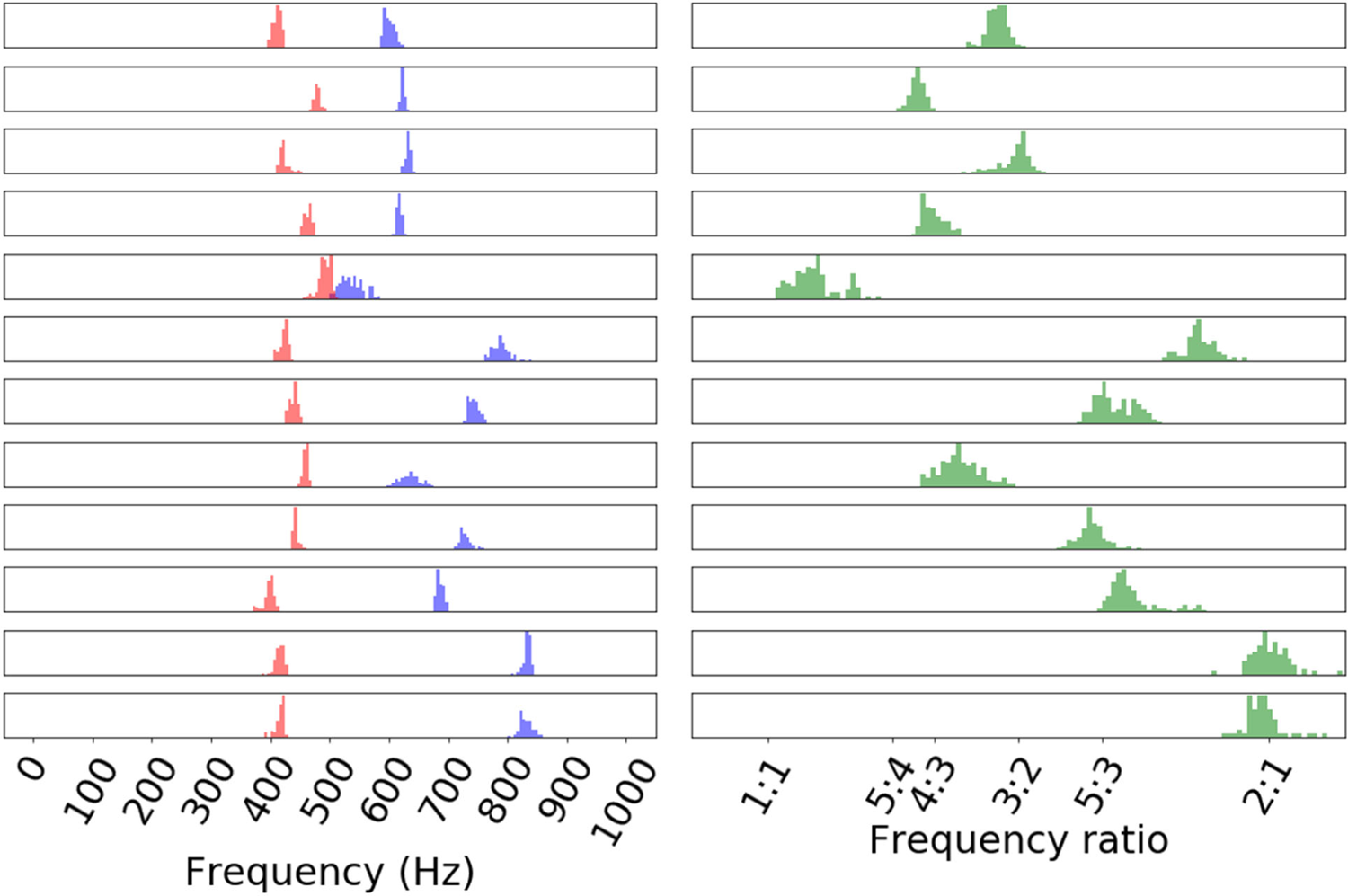
An example of the existence of ‘phonotypes’. Individual mosquitoes, and pairs of mosquitoes, occupy narrow ranges of frequencies, and frequency ratios. These narrow individual distributions introduce distinct peaks when used to calculate population histograms, which will only be averaged out by a sufficiently wide, and sufficiently representative, sampling of the underlying population (data from *Aldersley et al., 2016* (*22*)).

**Fig. S13.**
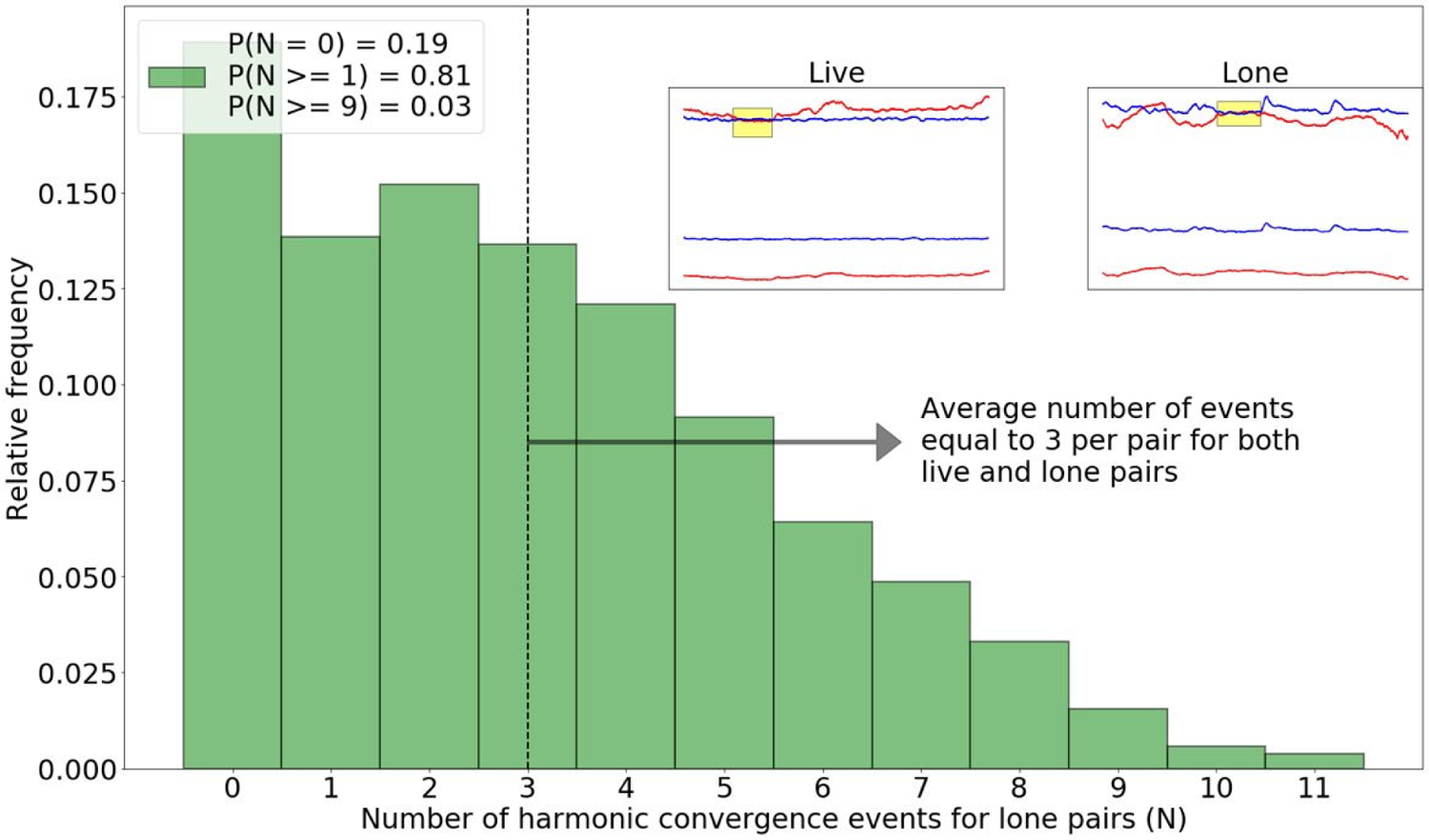
Relative frequency distribution of the number of harmonic convergence events (N) counted for the 513 lone (virtual) pairs. This distribution serves as reference (null hypothesis) to test if harmonic convergence events are significantly different in live (real) pairs. For an individual harmonic convergence event to be of statistical significance (i.e. to have a probability p<0.05 of having occurred by chance), a live pair must exhibit at least 9 harmonic convergence events during the one-minute long flight. Any value below 9 does not constitute a statistically noteworthy event (i.e. it has a probability p>0.05 of having occurred by chance). The average number of harmonic convergence events (see also Figs. 4 and S14) for both lone and live pairs is ∼ 3. The insets illustrate an example of harmonic convergence at 3:2 for live and lone pairs (blue: male; red: female; shown are fundamentals, the male’s 2^nd^ harmonic and the female’s 3^rd^ harmonic; harmonic convergence events highlighted in yellow).

**Fig. S14.**
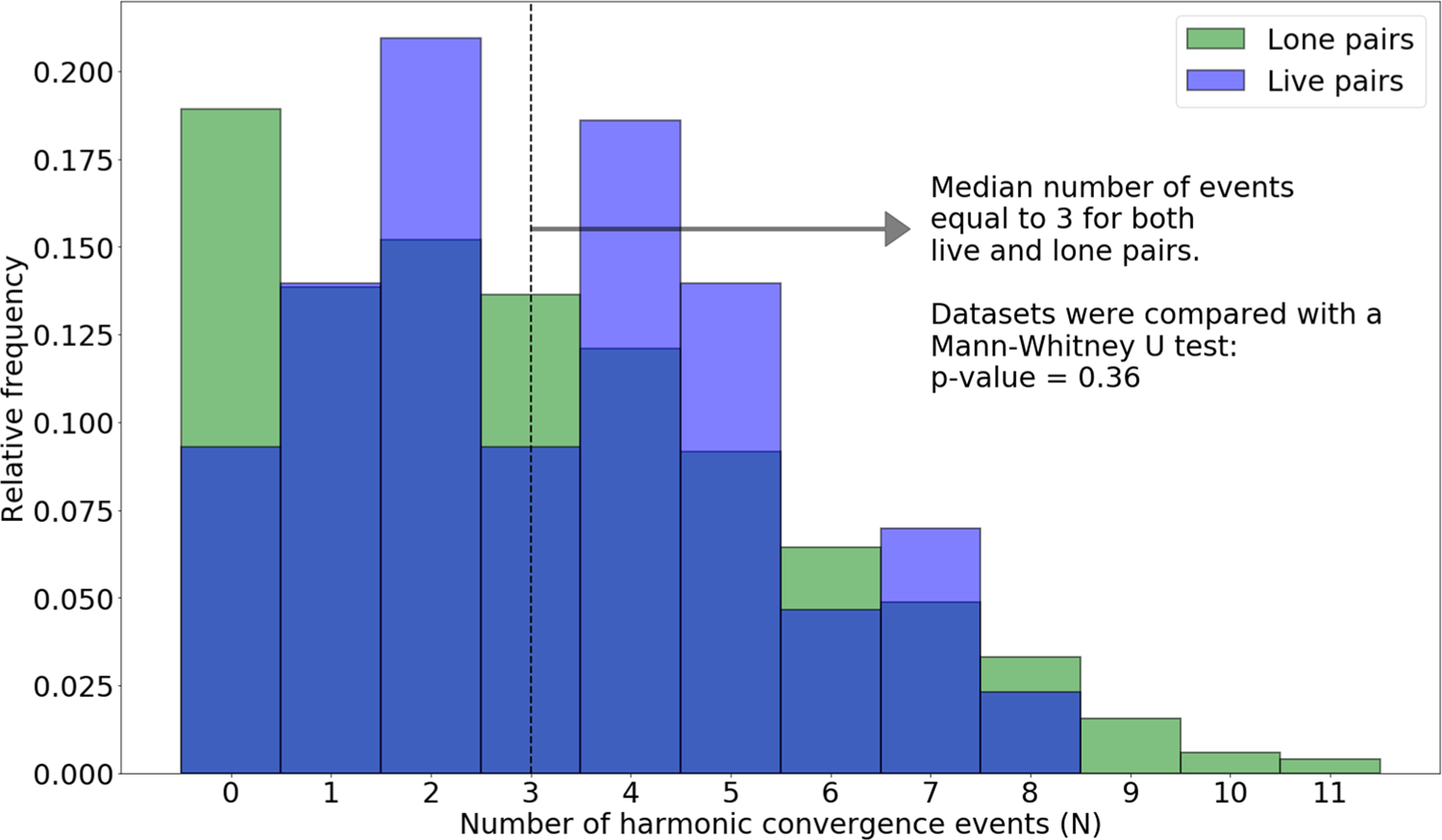
Superimposed relative frequency distributions of the 513 *lone* (green), and 43 *live* (blue) pairs of mosquitoes. Indicated in the figure is the result of the comparison of the two datasets that formed the distributions: Mann-Whitney U test with p-value = 0.36. A Mann-Whitney U test comparison of lone with playback gives a p-value of 0.17.

**Fig. S15.**
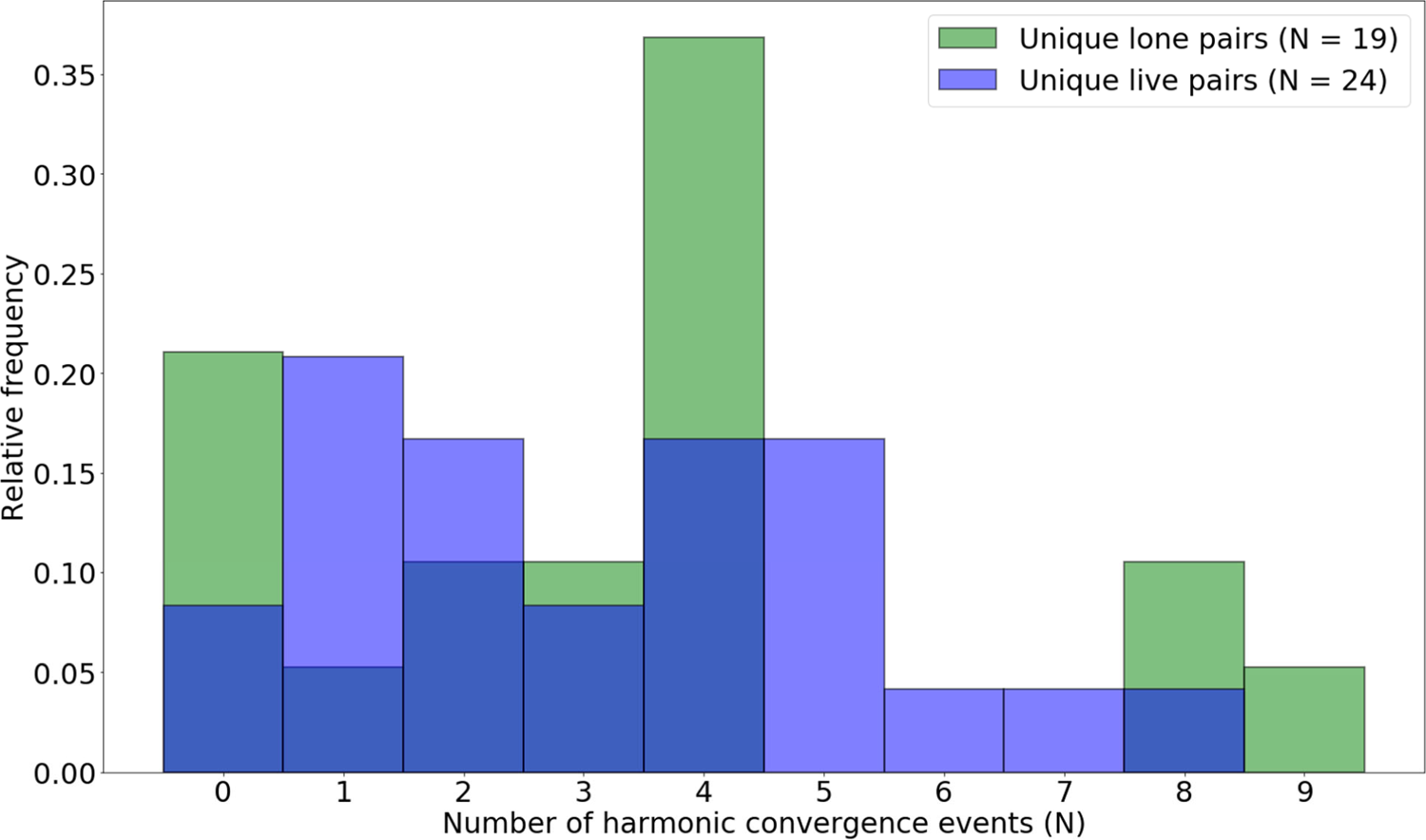
Superimposed relative frequency distributions of 19 unique *lone* (green), and 24 unique *live* (blue) pairs of mosquitoes. Comparison of the datasets with a Mann-Whitney U test gave a p- value of 0.98.

**Fig. S16.**
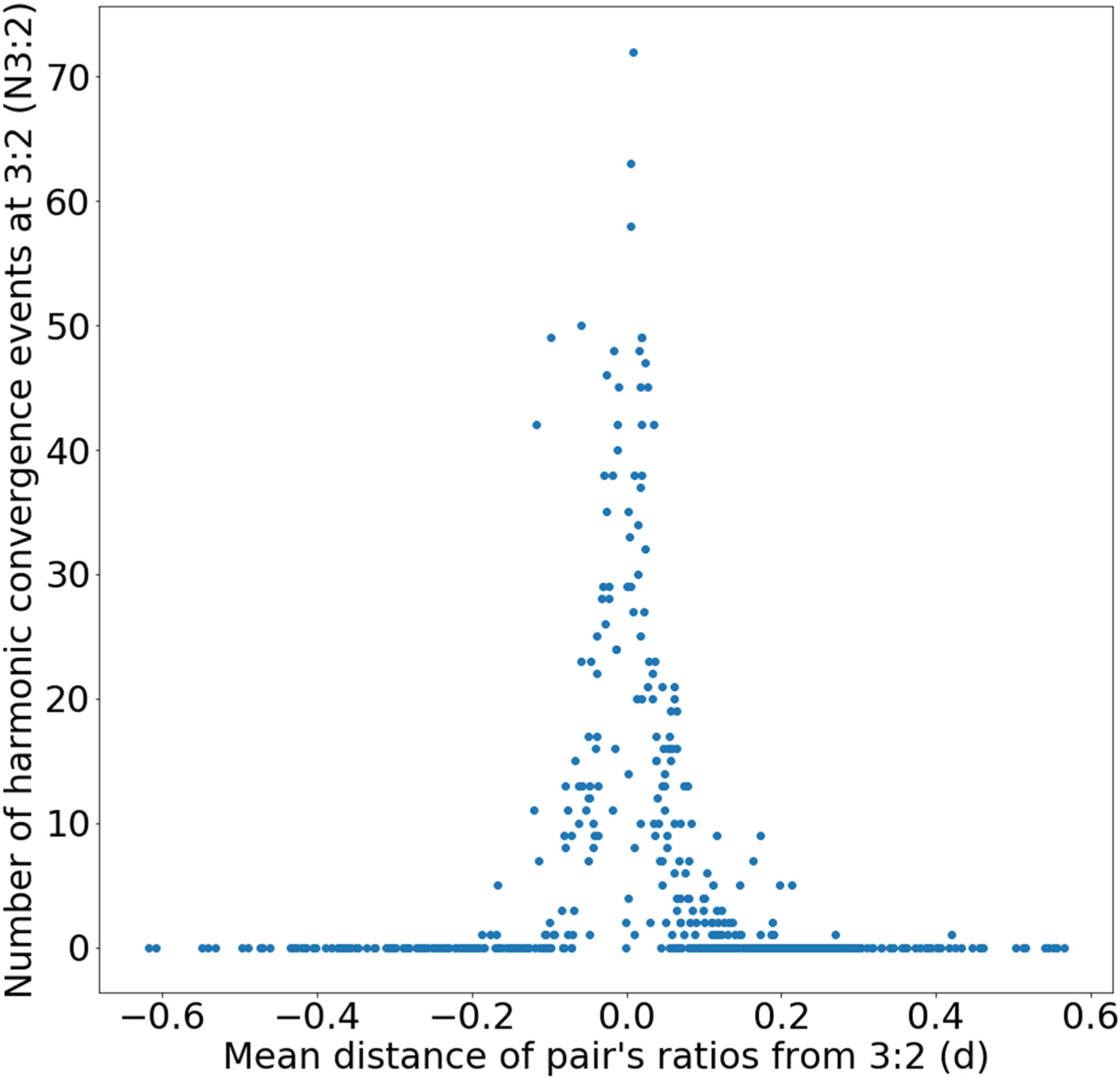
An illustration of how the number of harmonic convergence events at a given ratio (NHr) exhibited by a mosquito pair is a complex function of the mean distance (d) between the pairs’ mean frequency ratios and the given harmonic convergence ratio. Note that due to the 1s gap criterion the number of harmonic convergence events drops around zero distances (d=0). All of this introduces a very high sensitivity to noise around the given harmonic convergence ratio. Small random variations of recorded male or female flight tones (and resulting changes of the flight tone ratios) lead to dramatic increases or decreases in the number of harmonic convergence events, which can bias small sample size data sets.

### Annex 1: Probing harmonic convergence events for statistical signatures of acoustic interaction

#### 1.0 Synopsis

When flying in close proximity to each other, male and female mosquitoes have been suggested to interact acoustically by matching their flight tones, either at the level of the fundamental frequency (*34*) or the nearest shared harmonic (*21*). More precisely, for both *Aedes* (*21*), *Anopheles* (*35*) and *Culex* (*19*) mosquitoes, the 2^nd^ harmonic of the male flight tone (M2) was found to converge with the 3^rd^ harmonic of the female flight tone (F3) for short periods of time (<2s). Such a convergence event would correspond to a 3:2 (or 1.5) ratio of the corresponding fundamental flight tones.

We observed a daily shift of fundamental flight tones in male *Anopheles*. Notably, males were caged separately from their female counterparts and the respective cages kept in different incubators, thus ruling out any interactions between the sexes; the males’ flight tone shifts were not mirrored by their conspecific females. As a result of the males’ flight tone modulations, the corresponding male:female flight tone ratios moved closer to values of ∼1.5 during the daily activity peaks of *Anopheles* mosquitoes. We thus wondered if the described 3:2 convergence events simply reflected the random harmonic overlap produced by the males’ circadian maintenance of their fundamental flight tones. Rather than signaling an acoustic interaction between male and female, increases in harmonic convergence events would simply be the result of random fluctuations around a given pair’s median flight tone ratio, and its (likewise random) proximity to the respective harmonic convergence ratio (e.g. 1.5).

Robust tests of these relations are missing; only a single study (*22*) has explored this question experimentally before and in large enough sample size; they made their data publicly available at https://data.bris.ac.uk/data/dataset/1a44saul2ijj31u0raipvlims6. We conducted an in-depth statistical analysis of the study’s data set, and also applied some of the novel approaches introduced by the authors (*22, 36*). In a nutshell, we compared unique *real* (live) pairs to unique *virtual* (lone) pairs, making a deliberate attempt to reduce any distortions that arise from reusing individual pairs. We found that random overlaps are sufficient to explain both the spectral nature, and respective probability, of harmonic convergence events. An assumption of acoustic interaction is neither required nor statistically justified.

### 1.1 *Aedes aegypti* tethered flight data

The data set from *Aldersley et al.* (*2016*) (*22*), which we analyzed, used *Aedes aegypti* and tethered flight recordings throughout. Our own free-flight experiments confirm that just as in *Anopheles*, *Aedes* males – but not females – show daily (and state-dependent) modulations of their flight tones (Fig. S4 and S7), which include an upshift of flight tone frequencies during swarm time (dusk). Consistent with previous reports, the flight activities of *Aedes* mosquitoes (both males and females) were more evenly spread across the LD cycle and showed higher flight activities during the light phase than *Anopheles* (Fig. 2). Probably reflecting this enhanced level of behavioral activity, the male:female flight tone ratio in *Aedes* remained close to ∼1.5 for most parts of the day (Fig. S7, top). At swarm time (and driven by a male-specific increase in wing beat frequency) it rose further to values of ∼1.59 (Fig. S7, middle). Interestingly, though, analyses of the data from *Aldersley et al.* (*2016*) (*22*) reveal a flight tone ratio of only ∼1.4 during tethered flight (Fig. S7, bottom). Thus, just as in *Anopheles*, the flight tones of male *Aedes* are state-dependent and the corresponding (male:female) ratios remain close to the theoretical optimum of 1.5. We therefore scrutinized the data set made available by *Aldersley et al.* (*2016*) (*22*) to test if harmonic convergence events merely arose by chance.

The specific recordings used for each analysis are stated in the relevant sections. Our investigation focuses on two subsets of the authors’ database: (i) Flight tone recordings of *live* male-female pairs,(ii) *lone* males and females, and (iii) *live* male *playback* female ‘pairs’. A *live* (or real) pair is defined as a pair of tethered mosquitoes beating their wings in proximity to one another while the signal produced by each of their wingbeats (i.e. the flight tones) is recorded by separate microphones. These data are available at the above stated web resource under the heading ‘live_oppsex_pairs’. A lone male or female is defined as a tethered mosquito beating its wings in isolation (isolated from any other mosquito), while its flight tone is recorded by a microphone. Male and female *lone* recordings are also available at the web resource under the heading ‘lone_males’ and ‘lone_females’, respectively. A *lone* (or virtual) pair is produced by randomly pairing a lone male with a lone female. A *live* male *playback* female pair is defined as a lone tethered male beating its wings while being stimulated with the playback of a pre-recorded female (live or lone). These data are available at the above stated web resource under the heading ‘playback_pairs’. All flight tone recordings are one minute long and sampled at 40 kHz.

### 1.2 Flight tone extraction analysis

The flight tones of *Aedes aegypti* produced during tethered flight were extracted following a version of the protocol exemplified in *Aldersley et al. 2014* (*36*); here adapted to, and coded in, Python. Briefly: As an initial processing step, the recordings were bandpass filtered (lower cutoff point at 200Hz and higher cutoff point at 1,000Hz). The one-minute long recordings were segmented into non-overlapping windows of length *τ* = 100ms. Starting with the first segment (i.e. 0 - 100ms) a Fast Fourier Transform (FFT) was then applied to each segment. From the FFT we extracted *f*_L_ and *f*_H_, the distances (in Hz) to the left and right of *f*_o_, the most prominent peak in the frequency domain. Here *f*_L_ and *f*_H_ are given by:

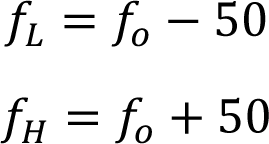

These two limits were then used to supply a bandwidth for bandpass-filtering the segment. The Hilbert transform was subsequently applied to the bandpass-filtered segment to extract the respective instantaneous frequency of the mosquito’s flight tone over that segment. This procedure was repeated for all flight segments, which were then appended together into the instantaneous frequencies of the complete 1-minute long flight. Finally, as described in *Aldersley et al., 2016* (*22*), the same piecewise aggregate approximation (PAA) averaging procedure, with window *w* = 0.5 s, was applied to the instantaneous frequencies to reduce noise. The result was a set of 120 frequency points (*w =* 0.5s, resulting in 2 frequency values per second over the entire 1 minute of tethered flight) for each mosquito recording.

### 1.3 Reconstruction of figure 6a from *Aldersley et al. 2016* (*22*)

Forty-three *live* pairs of male-female *Aedes aegypti* were originally used by the authors to generate a histogram of frequency ratios (*4*). Twenty-four of these forty-three pairs were unique pairs (i.e. different male *or* different female), with the remainder re-using individual pairs (i.e. same male *and* same female). For the construction of unique pairs, the original study by *Aldersley et al., 2016* (*22*) also repeatedly re-used individual males or females. For example: Unique pair *A* consists of male *m*_1_ and female *f*_1_ and unique pair *B* consists of male *m*_1_ and female *f*_2_. The frequency ratios of the forty-three live pairs were calculated here, in accordance with the authors analysis as follows:

A set of 120 frequencies, {*f*_1_, *f*_2_, …, *f*_120_}, was obtained for each mosquito of a pair by applying the above mentioned PAA averaging procedure over the minute-long flight. If we define *f*_mi,j_ and *f*_fi,j_ as the *i^th^* frequencies obtained from the male (*m*) and female (*f*) of the *j^th^* pair. Then the *i^th^* instantaneous frequency ratio of this pair is given by:

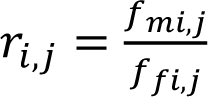

Resulting in a set of 120 instantaneous frequency ratios for each pair. The *r*_i,j_ were pooled together to reconstruct the frequency ratio histogram for live mosquito pairs (bin width = 0.01).

Sixty-eight (virtual) pairs, created from *lone* male and *lone* female *Aedes aegypti*, were used to create a frequency ratio histogram in the original analysis by *Aldersley et al., 2016* (*22*). These were constructed as follows: The study’s database contained data from 27 lone males and 19 lone females. These were combined by *Aldersley et al., 2016* (*22*) to create a total of 513 lone pairs, from which 68 were randomly sampled (the specific identity of these samples is unknown).

*Aldersley et al., 2016* (*22*) then conducted a frequency ratio analysis for these *lone* (virtual) pairs and compared them to the frequency ratio histogram from *live* (real) pairs. For our own analysis, all 513 lone pairs were used to reconstruct the complete frequency ratio histogram for *lone* (virtual) mosquito pairs (bin width same as above) (Fig. S9).

### 1.4 Deconstruction of Figure 6a from *Aldersley et al., 2016*

A key observation made by *Aldersley et al., 2016* (*22*) was the difference between (male:female) flight tone frequency ratio histograms from *live* (real) and *lone* (virtual) pairs. While lone pairs showed a smooth, (near unimodal) distribution, *live* (real) pairs showed a multimodal distribution, with clearly identifiable peaks at distinct frequency ratios. To test if these differences, rather than reflecting unique acoustic interactions within live pairs, are caused by the different sample sizes used (*live* n=43, *lone* n=68) pairs, we randomly selected 43 and 68 lone pairings from the total of 513 and used these subsets to produce frequency ratio histograms.

Doing so, illustrates how the histogram differences seen by *Aldersley et al., 2016* (*22*) between *live* and *lone* pairs (and also confirmed by our own analyses, see *Fig. S9*) can be qualitatively reproduced solely comparing two sets of *lone* pairs (with n=43 and n=68, respectively) (*Fig. S11*).

### 1.5 Probability distribution of the number of harmonic convergence events of lone pairs

We now probed if harmonic convergence events were more likely to occur in live pairs. *Aldersley et al., 2016* (*22*) defined a harmonic convergence event as a male:female frequency ratio *r*_i,j_ (defined above) that meets the following three criteria:

(i) The ratio falls within any of the following ranges: {[0.99 - 1.01], [1.2375 - 1.2625], [1.3199 - 1.3466], [1.485 - 1.515], [1.65 - 1.6833], [1.98 - 2.02]}; that is, it is within ±1% of any of six ratios {1/1, 5/4, 4/3, 3/2, 5/3, 2/1}. These are the ratios that would result from harmonic convergence occurring at the harmonic ratios (male:female) 1:1, 5:4, 4:3, 3:2, 5:3, 2:1.
(ii) The ratio is repeated such that it falls within the respective range for at least 1s (that is, it is repeated in at least two subsequent time windows, given the PAA window of 0.5 s). In other words, the pair maintains harmonic convergence for at least one second.
(iii) the ratio occurs at least 1s after the termination of a preceding harmonic convergence event. That is, there should be a gap of at least one second between any two harmonic convergence occurrences if the second occurrence is to qualify as a harmonic convergence event.

Following these criteria, we calculated the random variable *N*, defined as the number of harmonic convergence events produced by each of the 513 lone pairs. We then calculated the corresponding relative frequency distribution (Fig. S13). Given the large number of samples, this relative frequency distribution approximates the underlying probability distribution and thus served us as reference distribution for our null hypothesis, which would posit that all harmonic convergence events are the result of random frequency overlaps between male and female flight tones. Similarly, the number of harmonic convergence events produced by *live* pairs was determined and the corresponding averages (mean and median) were computed for comparison against the reference distribution (Fig. S14 shows results for all pairings, Fig. S15 for unique pairs only).

### 1.6 The number of harmonic convergence events as a function of the mean distance from the harmonic convergence ratio

Let us define *d*_j_ as the mean of the distances of a pair’s ratios from a particular harmonic ratio *Hr* such that:

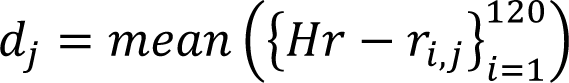

Where *Hr* can take the values {1, 5/4, 4/3, 3/2, 5/3, 2}. Focusing here on *Hr* = 3/2 = 1.5 (which is the originally proposed and most widely used harmonic convergence ratio (see ref. (*2*)), these distance metrics were calculated for live and lone pairs. In addition, the number of harmonic convergence occurrences at that ratio *j*, *N*_Hr,j_, were also counted for all live and lone pairs, chosen such that an occurrence meets criterion (i) of section 1.5. *N*_Hr_ was then plotted as a function of *d* for live and lone pairs (Fig. S16).

Criterion (i) for harmonic convergence of section 1.5 states that a harmonic convergence event occurs whenever the male:female frequency ratio *r*_i,j_ falls within ±1% of any of six ratios {1, 5/4, 4/3, 3/2, 5/3, 2} (though note that here we are only interested in ratio 3/2). Concentrating on this core HC criterion, provides a sense of how – for a given *Hr* - the amount of time a pair spends in a state of convergence is a simple function of the mean distance of the pair’s ratios from the respective *Hr* (here: 3/2). Importantly, this perspective has the advantage of being independent of how a harmonic convergence event has been defined by the experimenter in terms of time duration (Fig. S16).

### 1.7 Comparison of the distances of pair ratios from the harmonic convergence ratio: Live versus lone pairs

Let us define *d*_a,j_ as the mean of the absolute distances of a pair’s ratios from a harmonic ratio *Hr* (i.e. the variance of *r*_i,j_ about *Hr*) such that:

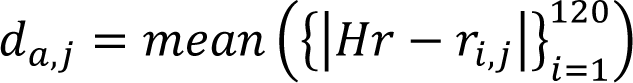

Where *Hr =* 1.5, as in section 1.6 This distance statistic was computed for each pair of the group of 24 unique live and the group of 19 unique lone pairs of mosquitoes. A comparison of the two groups (Table S1) was conducted using a Mann-Whitney U test (p-value = 0.93).

**Table S1.** Mean distance of pair ratios from the harmonic convergence ratio 3:2

|  |  |
| --- | --- |
| **Live (*Real*)** | **Lone (*Virtual*)** |
| 0.048 | 0.264 |
| 0.042 | 0.159 |
| 0.021 | 0.167 |
| 0.157 | 0.401 |
| 0.160 | 0.071 |
| 0.130 | 0.059 |
| 0.362 | 0.163 |
| 0.058 | 0.248 |
| 0.192 | 0.168 |
| 0.049 | 0.092 |
| 0.138 | 0.280 |
| 0.118 | 0.290 |
| 0.494 | 0.264 |
| 0.146 | 0.063 |
| 0.561 | 0.053 |
| 0.037 | 0.134 |
| 0.226 | 0.540 |
| 0.230 | 0.208 |
| 0.327 | 0.070 |
| 0.170 |  |
| 0.170 |  |
| 0.549 |  |
| 0.320 |  |
| 0.487 |  |

### 1.8 Comparison of flight tone distribution parameters: Live versus lone pair males

The flight tones of each of the 24 males and 24 females that were used in unique live pairs and 27 males and 19 females used in lone pairs were computed as exemplified in section 1.3.

Gaussian curves were then fitted on each frequency distribution, extracting the parameters of center, *μ,* and spread, *σ*, for each (Table S2). These statistics were subsequently collected for, and compared between, the live and lone groups using Mann-Whitney U tests. Specifically, four tests were conducted comparing: means (μ) between live males and lone males (p-value = 0.34), standard deviations (σ) between live males and lone males (p-value = 0.74), means between live females and lone females (p-value = 0.08), and standard deviations between live females and lone females (p-value = 0.50).

**Table S2.** Individual mosquito flight tone distribution Gaussian fit parameter extraction (μ=mean; σ = standard deviation)

| **Table S2:** Individual mosquito flight tone distribution Gaussian fit parameter extraction ($\mu$=mean; $\sigma$ = standard deviation) | | | | | | | |
| --- | --- | --- | --- | --- | --- | --- | --- |
| **Live male**<br>**$\mu$ (Hz)** | **Lone male**<br>**$\mu$ (Hz)** | **Live male**<br>**$\sigma$ (Hz)** | **Lone male**<br>**$\sigma$ (Hz)** | **Live female**<br>**$\mu$ (Hz)** | **Lone female**<br>**$\mu$ (Hz)** | **Live female**<br>**$\sigma$ (Hz)** | **Lone female**<br>**$\sigma$ (Hz)** |
| 691.5 | 629.19 | 23.6 | 9.9 | 465.26 | 509.18 | 19.59 | 11.47 |
| 600.28 | 692.52 | 8.43 | 8.56 | 411.82 | 488.7 | 6.78 | 7.65 |
| 632.32 | 806.49 | 4.24 | 8.09 | 423.37 | 448.76 | 8.76 | 7.68 |
| 628.13 | 658.0 | 3.51 | 14.21 | 467.92 | 489.94 | 7.96 | 3.24 |
| 593.06 | 482.69 | 4.93 | 3.78 | 442.83 | 465.01 | 7.75 | 15.63 |
| 666.05 | 641.96 | 69.98 | 8.15 | 448.13 | 439.35 | 7.21 | 4.12 |
| 787.84 | 605.08 | 13.91 | 6.06 | 423.24 | 452.53 | 7.11 | 7.37 |
| 696.97 | 646.27 | 8.62 | 3.88 | 447.41 | 459.73 | 6.53 | 57.58 |
| 742.39 | 671.84 | 8.89 | 16.2 | 438.96 | 517.53 | 6.81 | 23.21 |
| 696.91 | 633.08 | 23.9 | 12.21 | 470.28 | 463.44 | 6.94 | 7.87 |
| 709.12 | 641.96 | 26.94 | 8.15 | 432.92 | 503.64 | 6.39 | 18.1 |
| 635.2 | 745.72 | 16.06 | 10.56 | 459.55 | 460.15 | 3.79 | 25.02 |
| 488.97 | 666.38 | 8.55 | 73.42 | 486.13 | 462.51 | 12.21 | 14.53 |
| 728.58 | 714.65 | 10.12 | 11.36 | 442.62 | 408.71 | 4.1 | 3.1 |
| 647.14 | 566.52 | 19.89 | 4.1 | 689.14 | 424.51 | 8.57 | 11.7 |
| 675.69 | 559.19 | 7.14 | 50.23 | 440.12 | 433.45 | 4.33 | 4.3 |
| 686.49 | 628.76 | 5.51 | 14.91 | 397.96 | 483.97 | 9.25 | 3.91 |
| 675.14 | 861.76 | 6.61 | 11.2 | 390.59 | 512.57 | 13.09 | 22.54 |
| 779.56 | 723.57 | 12.83 | 10.85 | 427.58 | 505.6 | 18.05 | 6.32 |
| 869.13 | 783.56 | 13.33 | 11.94 | 520.85 |  | 14.83 |  |
| 860.44 | 670.57 | 9.99 | 8.98 | 515.28 |  | 7.75 |  |
| 852.44 | 618.07 | 26.28 | 14.86 | 416.06 |  | 8.57 |  |
| 652.54 | 865.51 | 11.94 | 13.19 | 552.99 |  | 7.48 |  |
| 685.71 | 642.07 | 9.19 | 21.56 | 345.29 |  | 7.44 |  |
|  | 836.94 |  | 10.64 |  |  |  |  |
|  | 671.13 |  | 6.6 |  |  |  |  |
|  | 666.36 |  | 8.02 |  |  |  |  |

### 1.9 Comparison of flight tone diversities: Live versus lone pair males

The *uncertainty* (or it’s inverse, the *information*) inherent to a random variable, e.g. the male mosquito’s flight tone *f*_m_ can be quantified using a measure such as the *Shannon Entropy* of *f*_m_. The Shannon Entropy (also known as Shannon’s Diversity Index) (*37*) can be restated, in the context of mosquito flight tones, as a measure of how many frequency channels the mosquito occupies and how often it occupies each of these during its flight. In other words, it is a metric of how diverse a mosquito’s landscape of flight tones is. Here the Shannon Entropies of the flight tones of the 24 males and 24 females that were used in unique live pairs and the 27 males and 19 females used in unique lone pairs were computed as follows:

First, the set of each mosquito’s flight tones were discretized, and assigned to bins of 5Hz width along the mosquito’s flight tone range. Then, the proportion *p*_rb_ of the number of a male’s flight tones *n* falling in bin *b* relative to the total number of flight tones is given by:

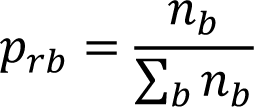

Then the Shannon entropy *H* of a mosquito’s flight tones is defined us:

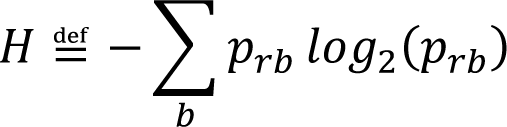

The Shannon Entropies of live males’ flight tones were compared to those of lone males’ flight tones using a t-test (p-values: 0.41, 0.73 for male and female comparisons respectively).

**Table S3.** Mosquito flight tone diversities (Shannon)

| **Table S3: Mosquito flight tone diversities (Shannon)** |  |  |  |
| --- | --- | --- | --- |
| **Live male** | **Lone male** | **Live female** | **Lone female** |
| 3.884 | 2.839 | 3.671 | 2.928 |
| 2.686 | 2.665 | 2.427 | 2.642 |
| 1.821 | 2.659 | 2.412 | 2.449 |
| 1.618 | 3.375 | 2.672 | 1.523 |
| 2.022 | 1.793 | 2.614 | 3.45 |
| 5.075 | 2.735 | 2.473 | 1.801 |
| 3.358 | 2.325 | 2.41 | 2.314 |
| 2.791 | 1.664 | 2.41 | 3.229 |
| 2.79 | 3.424 | 2.466 | 4.01 |
| 3.911 | 2.618 | 2.477 | 2.664 |
| 3.944 | 2.735 | 2.415 | 3.198 |
| 3.641 | 2.815 | 1.735 | 2.736 |
| 2.786 | 4.149 | 3.02 | 3.437 |
| 2.85 | 2.932 | 1.646 | 1.494 |
| 3.143 | 1.859 | 2.813 | 2.851 |
| 2.535 | 3.023 | 1.997 | 1.885 |
| 2.157 | 3.457 | 2.677 | 1.711 |
| 2.333 | 3.088 | 2.738 | 3.365 |
| 3.271 | 2.957 | 3.412 | 2.25 |
| 3.151 | 3.178 | 3.341 |  |
| 2.92 | 2.716 | 2.608 |  |
| 4.084 | 3.303 | 2.745 |  |
| 2.945 | 3.261 | 2.58 |  |
| 2.741 | 3.631 | 1.83 |  |
|  | 3.026 |  |  |
|  | 2.376 |  |  |
|  | 2.651 |  |  |

## References and Notes

1. X. Y. Li, H. Kokko, Sex-biased dispersal: a review of the theory. Biol. Rev. 94, 721–736 (2019).

2. W. Takken et al., in Bridging Laboratory and Field Research for Genetic Control of Disease Vectors, B. G. J. Knols, C. Louis, Eds. (2006), vol. 11, pp. 183-+.

3. P. I. Howell, B. G. J. Knols, Male mating biology. Malaria Journal 8, 10 (2009).

4. J. D. Charlwood, M. D. R. Jones, Mating in the mosquito, Anopheles gambiae s.l. Physiological Entomology 5, 315–320 (1980).

5. A. Diabate et al., Spatial distribution and male mating success of Anopheles gambiae swarms. Bmc Evolutionary Biology 11, (2011).

6. S. B. Poda, et al., Sex aggregation and species segregation cues in swarming mosquitoes: role of ground visual markers. Parasite Vector 12, (2019).

7. S. P. Sawadogo et al., Targeting male mosquito swarms to control malaria vector density. Plos One 12, (2017).

8. E. W. Kaindoa et al., Swarms of the malaria vector Anopheles funestus in Tanzania. Malaria Journal 18, (2019).

9. J. D. Charlwood, R. Thompson, H. Madsen, Observations on the swarming and mating behaviour of Anopheles funestus from southern Mozambique. Malaria Journal 2, (2003).

10. C. Montell, L. J. Zwiebel, in Progress in Mosquito Research, A. S. Raikhel, Ed. (Academic Press Ltd-Elsevier Science Ltd, London, 2016), vol. 51, pp. 293-328.

11. C. Johnston, Auditory apparatus of the Culex mosquito. Q. J.Microsc. Sci. 3, 97–102 (1855).

12. M. C. Göpfert, D. Robert, Active auditory mechanics in mosquitoes. Proceedings of the Royal Society B: Biological Sciences 268, 333–339 (2001).

13. Joerg T. Albert, Andrei S. Kozlov, Comparative Aspects of Hearing in Vertebrates and Insects with Antennal Ears. Current Biology 26, R1050–R1061 (2016).

14. M. C. Göpfert, H. Briegel, D. Robert, Mosquito hearing: Sound-induced antennal vibrations in male and female *Aedes aegypti*. Journal of Experimental Biology 202, 2727–2738 (1999).

15. M. C. Göpfert, D. Robert, Nanometre-range acoustic sensitivity in male and female mosquitoes. Proc. R. Soc. Lond. Ser. B-Biol. Sci. 267, 453–457 (2000).

16. M. P. Su, M. Andrés, N. Boyd-Gibbins, J. Somers, J. T. Albert, Sex and species specific hearing mechanisms in mosquito flagellar ears. Nature Communications 9, 3911 (2018).

17. G. Gibson, B. Warren, I. J. Russell, Humming in Tune: Sex and Species Recognition by Mosquitoes on the Wing. Jaro-Journal of the Association for Research in Otolaryngology 11, 527–540 (2010).

18. F. Julicher, D. Andor, T. Duke, Physical basis of two-tone interference in hearing. P Natl Acad Sci USA 98, 9080–9085 (2001).

19. B. Warren, G. Gibson, I. J. Russell, Sex Recognition through Midflight Mating Duets in Culex Mosquitoes Is Mediated by Acoustic Distortion. Current Biology 19, 485–491 (2009).

20. P. M. V. Simões, R. Ingham, G. Gibson, I. J. Russell, Masking of an auditory behaviour reveals how male mosquitoes use distortion to detect females. Proc. R. Soc. B 285, 20171862 (2018).

21. L. J. Cator, B. J. Arthur, L. C. Harrington, R. R. Hoy, Harmonic convergence in the love songs of the dengue vector mosquito. Science 323, 1077–1079 (2009).

22. A. Aldersley, A. Champneys, M. Homer, D. Robert, Quantitative analysis of harmonic convergence in mosquito auditory interactions. Journal of the Royal Society Interface 13, (2016).

23. G. Wang et al., Clock genes and environmental cues coordinate *Anopheles* pheromone synthesis, swarming, and mating. Science 371, 411–415 (2021).

24. S. M. Villarreal, O. Winokur, L. Harrington, The Impact of Temperature and Body Size on Fundamental Flight Tone Variation in the Mosquito Vector Aedes aegypti (Diptera: Culicidae): Implications for Acoustic Lures. Journal of Medical Entomology 54, 1116–1121 (2017).

25. T. Nakata et al., Aerodynamic imaging by mosquitoes inspires a surface detector for autonomous flying vehicles. Science 368, 634-+ (2020).

26. P. M. V. Simões, R. A. Ingham, G. Gibson, I. J. Russell, A role for acoustic distortion in novel rapid frequency modulation behaviour in free-flying male mosquitoes. Journal of Experimental Biology 219, 2039–2047 (2016).

27. D. N. Lapshin, D. D. Vorontsov, Frequency tuning of swarming male mosquitoes (Aedes communis, Culicidae) and its neural mechanisms. Journal of Insect Physiology 132, 104233 (2021).

28. M. P. Su, et al., Assessing the acoustic behaviour of Anopheles gambiae (s.l.) dsxF mutants: implications for vector control. Parasite Vector 13, (2020).

29. A. N. Clements, The biology of mosquitoes. Vol. 2: Sensory reception and behaviour. (CABI Publishing, New York, 1999).

30. L. O. Araripe, J. R. A. Bezerra, G. B. d. S. Rivas, R. V. Bruno, Locomotor activity in males of Aedes aegypti can shift in response to females’ presence. Parasite Vector 11, 254 (2018).

31. Z. W. Dou et al., Acoustotactic response of mosquitoes in untethered flight to incidental sound. Scientific Reports 11, (2021).

32. R. J. Bomphrey, T. Nakata, N. Phillips, S. M. Walker, Smart wing rotation and trailing- edge vortices enable high frequency mosquito flight. Nature 544, 92 (2017).

33. Q. Geissmann, L. Garcia Rodriguez, E. J. Beckwith, G. F. Gilestro, Rethomics: An R framework to analyse high-throughput behavioural data. PLOS ONE 14, e0209331 (2019).

34. G. Gibson, I. Russell, Flying in tune: Sexual recognition in mosquitoes. Current Biology 16, 1311–1316 (2006).

35. C. Pennetier, B. Warren, K. R. Dabiré, I. J. Russell, G. Gibson, “Singing on the Wing” as a Mechanism for Species Recognition in the Malarial Mosquito Anopheles gambiae. Current Biology 20, 131–136 (2010).

36. A. Aldersley, A. Champneys, M. Homer, D. Robert, Time-frequency composition of mosquito flight tones obtained using Hilbert spectral analysis. J Acoust Soc Am 136, 1982–1989 (2014).

37. C. E. Shannon, W. Weaver, The mathematical theory of communication. (University of Illinois Press, 1949).

